# The anoxic switch: Strain variation in H_2_ exchange during repeated illumination after dark anoxia

**DOI:** 10.64898/2026.10.28.762455

**Authors:** Michael Kikoine, Iftach Yacoby

## Abstract

Hydrogen (H_2_) exchange during transitions from dark incubation to sustained illumination provides a readout of physiological history in green microalgae. In *Chlamydomonas reinhardtii*, the first illumination after prolonged dark anoxia gives a higher H_2_ production rate than later illuminations, an “anoxic switch” linked to photosystem II (PSII) photosynthetic control. Membrane-inlet mass spectrometry was used to compare 54 independently grown cultures from 17 strains across sequential irradiances and 48 cultures from 15 strains during seven repeated cycles of 2 min of light and 3 min of darkness at strain-specific irradiances. For the primary early-response 30-s endpoint, first sustained-light rates exceeded the mean of cycles 5–7 in 43 of 48 cultures. Ratios of strain mean first to later rates exceeded 1 in 14 of 15 strains and ranged from 0.63 to 17.48, and strain accounted for most of the variance in culture-level ratios. G1122 was the only strain with higher later rates. The predominance of higher first rates persisted across alternative rate windows and after excluding cultures with elevated pre-light O_2_, although exact magnitudes depended on the window. O_2_ and nominal CO_2_ signal changes varied among strains, without a detected cross-strain correlation with the H_2_ ratio. An enhanced first-light H_2_ response was therefore the predominant pattern in strains of the Chlorellaceae, Scenedesmaceae, Selenastraceae and Chlamydomonadales, but its magnitude differed by more than an order of magnitude, identifying candidate strains for testing whether PSII photosynthetic control explains this variation.

## Introduction

The transition from dark anoxia to light requires green microalgae to restart photosynthetic electron transport at a time when the supply of electron acceptors is also changing. H_2_ metabolism is one way the cells manage this transition. Gaffron and Rubin (1942) first described fermentative and light-driven H_2_ production following anaerobic incubation. In *Chlamydomonas reinhardtii*, a chloroplast enzyme called hydrogenase takes electrons from ferredoxin and uses them to turn protons into H_2_. Mutants lacking hydrogenase show that this activity helps activate photosynthesis under anoxia (Ghysels et al., 2013).

The brief burst of H_2_ production reflects both how electrons are being routed in the cell and how much active hydrogenase is available. In *C. reinhardtii*, hydrogenase activity and cyclic electron flow contribute one after another to switching on carbon fixation (Godaux et al., 2015). Milrad et al. (2018) showed that H_2_ production can stop even before O_2_ inactivates the hydrogenase itself, because carbon fixation starts competing for the same electrons. A separate pathway, flavodiiron-driven O_2_ reduction, also helps switch on photosynthesis under anoxia (Burlacot et al., 2018). Measuring several dissolved gases together can therefore describe this transition, but pinning down which specific pathway is responsible needs further evidence.

The very first light exposure after a long period of darkness can behave differently from later ones. Milrad et al. (2018) reported that the initial H_2_ rate in *C. reinhardtii* was about three times higher than in later light–dark cycles. Milrad et al. (2023) then linked this drop-off over repeated illumination to regulation on the acceptor side of photosystem II (PSII), based on spectroscopic and biochemical measurements; a longer dark period restored the strong initial response. Alternating brief light and dark intervals can sustain efficient H_2_ photoproduction (Kosourov et al., 2018), indicating practical relevance for the dependence on illumination history. These findings raise a natural question: how consistent is this first-light enhancement across different green microalgal strains, and how much does its size vary?

Earlier screening already showed that H_2_ production varies among green algal lineages, and even among strains assigned to the same species (Timmins et al., 2009). Here we compare the first light exposure with later ones in a collection of G-coded strains, using membrane-inlet mass spectrometry (MIMS). We call this contrast, measured after an initial period of dark anoxia, the “anoxic-switch response”. We measure H_2_ responses across a range of light intensities, compare the first and later light periods within each culture, and track the accompanying O_2_ and CO_2_ changes. We ask how widespread this response is among green microalgae, how much its magnitude varies among strains, and which physiological explanations are consistent with the accompanying gas exchange.

## Materials and Methods

### Algal strains and identification

Strains are designated by collection codes independently of their working taxonomic names (Table S1). Shared identification and isolation records follow the companion characterization manuscript (Kikoine et al., n.d.). Morphology and ribosomal DNA provide supporting evidence rather than definitive species identification. G1099, G1100, G1106 and G1122 lack reliable molecular assignments and retain morphology-based names. The companion reanalysis also treats G1095, G1101, G1104 and G1121 as provisional and leaves the generic placement of G1105 unresolved. G1098 is the *C. reinhardtii* CC124 reference strain. G1093 is the laboratory *C. ohadii* culture linked by provenance to SAG 2657; its ribosomal marker does not independently distinguish that name from *C. sorokiniana*. G1102 was supplied as *Acutodesmus obliquus* SAG 276-3d; this name is synonymous with *Tetradesmus obliquus*, used for the separate culture G1097. Recorded provenance is summarized for the full collection in Table S9. Axenicity was not tested for the characterized collection; visual selection established unialgal cultures, and associated microorganisms may contribute to gas exchange.

### Cultivation and sample preparation

Cultures were maintained at 25 °C under continuous illumination (150 µmol photons m⁻² s⁻¹), with shaking under ambient air. Cultivation used the same vitamin stock composition as the companion characterization; the stock recipe is given in the supplementary Methods. Each biological replicate was an independently grown culture. Chlorophyll was quantified by DMSO extraction at 65 °C and absorbance at 665, 649 and 730 nm using Wellburn’s (1994) equations. Cell suspensions were assayed in TAP medium at 20 µg mL⁻¹ total chlorophyll. The extraction calculation and reference TAP recipe are given in the supplementary Methods. The FireSting cultivation atmosphere and assay settings in the companion paper are separate from the MIMS protocol.

### Calibration and reconstruction of gas concentrations

Two MIMS systems were used, with instrument- and filament-specific calibration profiles. H_2_ and O_2_ ion currents were converted using C(t) = mI(t) + b with the recorded coefficients and one application of background offsets (Tables S2 and S3). Original calibrated records were retained for rate calculations and operational O_2_-threshold assessments; neither gas was forced to zero for these analyses. Display-only shifts are described separately. CO_2_ was converted using its recorded affine coefficients but lacked independent absolute-concentration validation; its plotted scale and rates are therefore reported as nominal signal units and nominal units min⁻¹. Positive and negative signs describe changes on that scale. No argon normalization or gas-specific membrane-withdrawal correction was applied. All slopes describe net dissolved-pool or signal changes, including biological production/consumption and physical measurement effects; they are not gross biological fluxes.

### Gas measurements

A 5-mL suspension was measured in a sealed quartz cuvette connected to a home-built MIMS system based on the arrangement described by Liran et al. (2016), with continuous magnetic stirring, no headspace and no bubbling. Measurements were made at room temperature. A Dual-PAM-100 instrument (Walz) supplied illumination. Ion currents for masses 2, 32 and 44 (H_2_, O_2_ and CO_2_) were recorded as ASCII files; within-recording median gas-channel sampling intervals ranged from 2.124 to 2.599 s. Samples underwent an initial 1-h dark incubation before the measurement sequence. For this study, operational anoxia was defined as reconstructed bulk dissolved O_2_ below 10 µM. This pragmatic cutoff was used for classification and sensitivity analysis; it was not an experimentally established species-specific hydrogenase-inactivation threshold. Illuminations that started above this threshold were retained and examined by sensitivity analysis.

### Irradiance-response measurements

A sequential irradiance-response protocol characterized H_2_ exchange across the tested light intensities. Three 2-min conditioning light periods at 203 µmol photons m⁻² s⁻¹ were separated by 3-min dark intervals, following the timing used by Milrad et al. (2018). After the third conditioning period, a 5-min dark interval preceded six 2-min light periods at 52, 107, 203, 357, 722 and 1110 µmol photons m⁻² s⁻¹, applied in increasing order and separated by 3-min dark intervals (Fig. S1). H_2_, O_2_ and CO_2_ rate curves are shown in Figures S5–S7, respectively. Conditioning periods were excluded from the six-point curves. The peak irradiance was defined as the tested intensity with the highest mean peak H_2_ rate across biological replicates; it was a sampled maximum rather than a fitted optimum. Because irradiance increased with elapsed time and prior light exposure, the curves describe responses to this sequence rather than an isolated irradiance effect. Table S3 lists these sampled maxima and the irradiances used for repeated-cycle experiments.

### Repeated light-dark experiments

The repeated-cycle protocol comprised seven sustained 2-min light periods separated by 3-min dark intervals, with constant irradiance within each recording. Time zero was the onset of the first sustained measurement light period in the analyzed sequence; subsequent periods began at 5, 10, 15, 20, 25 and 30 min. Throughout this work, first illumination refers to this first 2-min measurement period. Phase timing was obtained from the recording metadata. Light fits excluded the exact phase endpoint; primary H_2_ fits ended before 120 s.

One repeated-cycle experimental group was retained per strain. Where matching recordings were available, the irradiance was selected to match the sampled H_2_ maximum in that strain’s irradiance-response series. Table S3 lists irradiances and biological replicate counts, and Figure S2 illustrates the protocol. G1106 and G1121 contributed irradiance-response data only; the repeated-cycle dataset therefore covers 15 strains. Rate estimation and first-to-late comparisons

Our main first-versus-later comparisons used an early-response 30-second rate calculation (Figures 1 and 2 and Figures S3 and S4). We analyzed each gas using its original timestamps and calibrated concentrations. Within the first 72 s of each phase, we located either a minimum (before an expected rise) or a maximum (before an expected fall) in the raw data. Starting 0– 18 s after this point, in 0.6-s steps, we tried candidate 30-second linear fits, staying within the first 120 s of the phase. We then picked the most positive slope for an expected rise, or the most negative slope for an expected fall. If every candidate slope had the wrong sign, we kept the one closest to the expected direction. H_2_ and O_2_ were expected to rise in light and fall in darkness; for CO_2_ it was the reverse. H_2_ and O_2_ rates are given in µM min⁻¹; CO_2_ uses nominal conversion-scale units as described above. Locating this raw minimum or maximum is simply a reproducible way to pick a fitting window — it does not mark a true physiological starting point. The supplementary Methods give the data-coverage criteria we required for a fit to count. For each culture, the first-light rate was paired with the mean light rate in cycles 5–7. Cultures received equal weight in strain means. Ratios in Figures 1 and 2 and Figures S3 and S4 divide the mean first-light rate by the mean later-light rate for each strain (the anoxic-switch ratio) and are shown only when both means have the expected sign. Figure 2 also reports H_2_ rate means ± sample standard deviation (SD). The number of cultures with a higher first than later rate was calculated from Table S4. Figure S2 applies the same window-selection procedure to selected individual recordings. No detection-limit threshold was applied, so ratios with near-zero denominators should be interpreted with caution.

**Figure 1:**
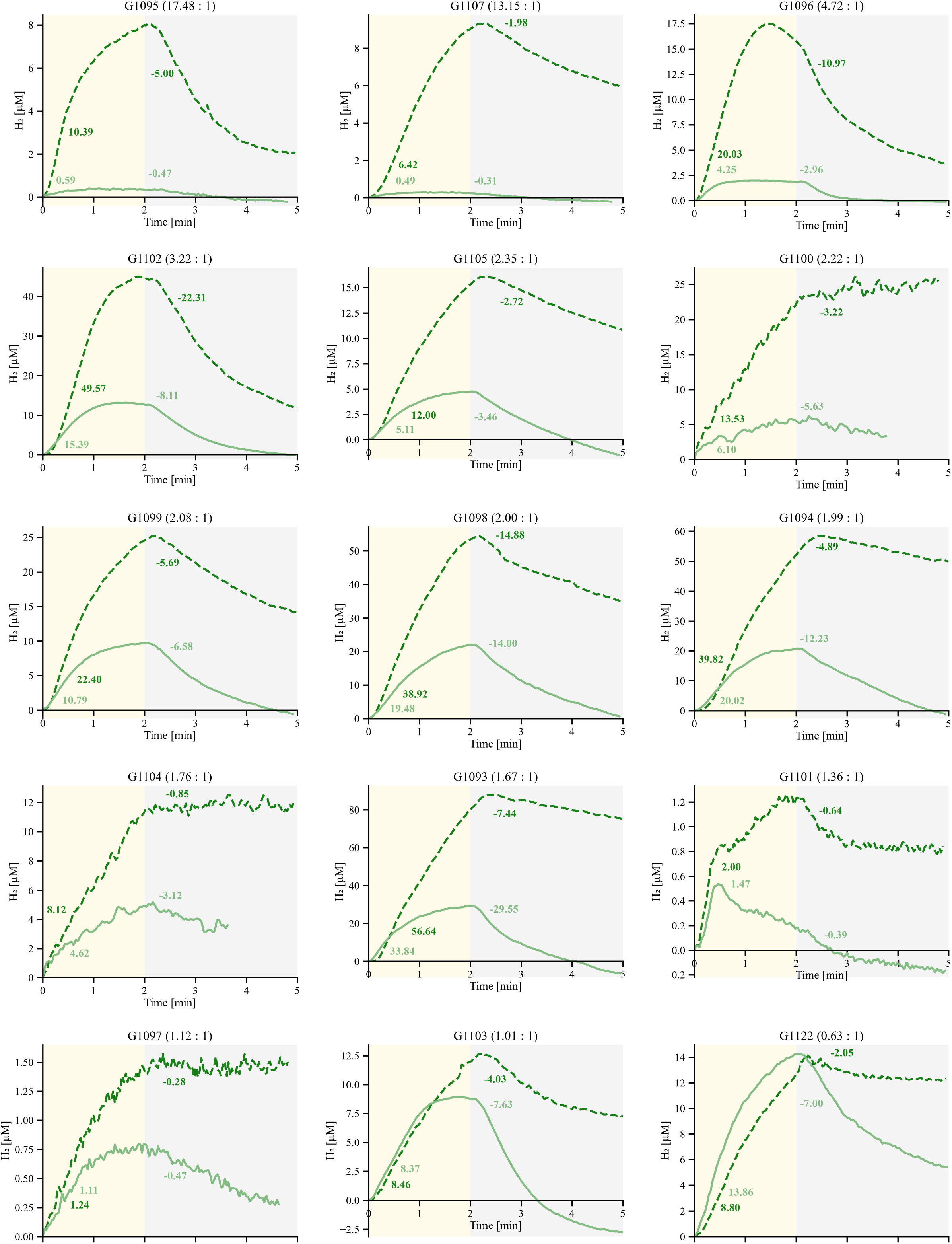
First and later H_2_ responses during repeated illumination. Each panel shows one of 15 strains (48 independently grown cultures in total; irradiances and n are given in Table S3). The dark-green dashed line is the mean first exposure; the light-green solid line is the mean of exposures 5–7, averaged first within each culture and then across cultures. We shifted each exposure to a common time-and-H_2_-concentration origin, so the curves show change from an exposure-specific baseline rather than absolute concentration. The yellow and gray backgrounds mark the nominal 2-min light / 3-min dark protocol after this alignment, so they do not mark the exact lamp transition on every trace. The numbers give the mean light and dark rates (µM min⁻¹) from the early-extremum 30-s method. The ratio in each panel heading is the mean first-light rate divided by the mean later-light rate; panels are ordered from highest to lowest ratio. Curves show means only, without error bands. Figure 2 and Table S4 use the same early-response estimates; Figure S8 shows a separate full-period calculation. First denotes the first sustained 2-min measurement light period in the analyzed sequence.

**Figure 2:**
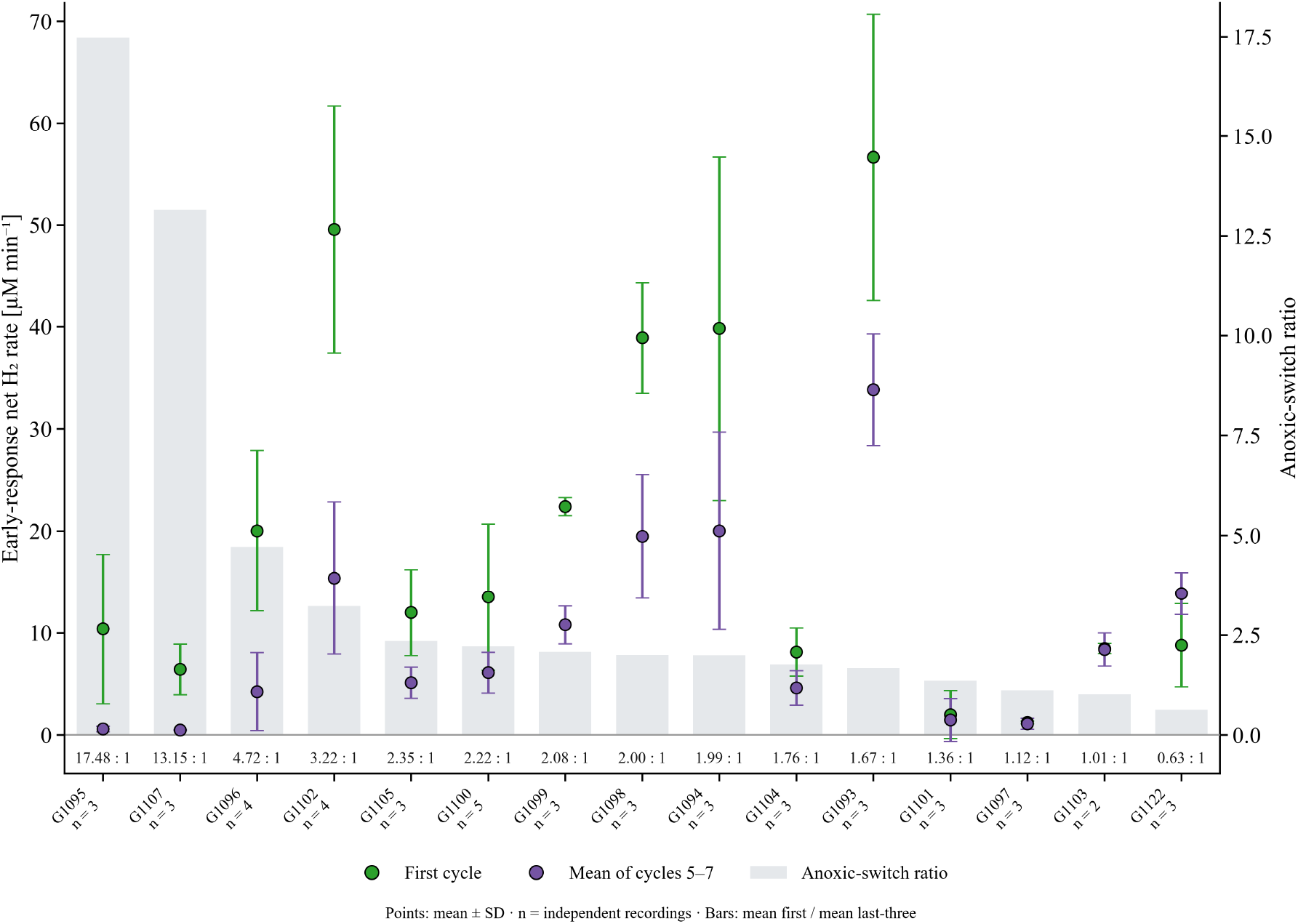
First and later H_2_ rates and their ratios. Each point is a strain mean ± SD, for the first light period (green) and for cycles 5–7 (purple; left axis). Later rates were first averaged within each culture; n gives the number of independent cultures (48 across 15 strains). The gray bars and values show the anoxic-switch ratio, mean first/mean later (right axis), with strains ordered from the highest to the lowest ratio. The early-response 30-s method here matches the one used in Figure 1. Tables S3 and S4 list the irradiances and the paired culture values. First denotes the first sustained 2-min measurement light period in the analyzed sequence.

A separate, full-period search characterized the irradiance-response curves and all seven cycles (Figures S5–S10). This search checked every eligible 30-s window across each full 2-min light period and the first 2 min of the following dark interval. We gave priority to the steepest slope in the expected direction; if none was found, we kept the strongest slope in the opposite direction, and a completely flat signal was recorded as zero. For the H_2_ sensitivity summaries, we divided each culture’s first rate by its mean of cycles 5–7 when both were positive, and summarized each strain by the median and range of these ratios. This differs from our main comparison in both how the window was chosen and how the result was summarized. Where the data allowed, we also calculated fixed-first-30-s and complete-120-s slopes. The supplementary Methods give the full details of these procedures, and the supplementary Results report the full-period comparison.

### Data presentation and O_2_ status

Recordings from independently grown cultures were treated as biological replicates. Irradiance-response and cycle-rate plots show individual cultures with arithmetic means ± standard error of the mean (SEM; Figs S5–S10). Concentration traces in Figure 1 and Figures S3 and S4 show means without error bands: cycles 5–7 were first averaged within each culture, followed by an equally weighted average across cultures. Each exposure was shifted in time and concentration to a detected local minimum for H_2_/O_2_ or maximum for CO_2_; a documented default onset was used when no qualifying extremum was detected. Aligned traces were interpolated onto a 1-s grid only within the measured range, whereas fitted rates used original timestamps and unaligned signals. Background shading denotes nominal protocol timing and does not identify the exact lamp transition on every aligned trace. Figures S1 and S2 show individual recordings shifted once to their H_2_ value at first illumination; these display adjustments did not alter calibration, fitted rates or O_2_ classifications. The sample size n denotes biological replicates; repeated cycles are observations within the same culture. Figure 2 summarizes paired H_2_ rates without time alignment, with SD bars showing variation among cultures.

For each illumination, we recorded the median O_2_ concentration over the preceding 30 s. The full-period analysis also recorded the maximum O_2_ concentration within its selected 30-s H_2_ window. We compared both values against our 10-µM operational anoxia threshold. Brief excursions above the cutoff that returned below 10 µM during the following dark interval were considered compatible with the repeated-cycle protocol and were not grounds for exclusion. Cycles that did not recover below the cutoff were also retained in the primary cohort, but their elevated pre-light O_2_ values were flagged and examined by sensitivity analysis. This classification used the original calibrated O_2_ signal, not the shifted display traces, so it depends on the absolute calibration.

Exploratory strain comparisons used natural-log culture-level first-to-later H_2_ ratios, with independently grown cultures as units and strain as a one-way factor. Ordinary ANOVA was accompanied by Welch’s unequal-variance ANOVA (Welch, 1951) and a median-centered Brown–Forsythe variance check. Games–Howell comparisons allowed unequal strain variances (Games & Howell, 1976). Their small-sample error control remains uncertain, so all pairwise results are exploratory. A sensitivity omitted G1095, G1096 and G1107. The share of variance associated with strain was summarized by η² and a one-way random-effects intraclass correlation. Cross-gas tests used strain ratios of means, permutation Spearman correlations and Holm adjustment across two comparisons (Tables S6–S8).

Sensitivity analyses retained the primary cohort and did not change the selected early-response rates. Fixed-first-30-s, whole-120-s and full-period peak-30-s slopes were compared using the same endpoint: the equally weighted strain mean first rate divided by the strain mean of within-culture rates in cycles 5–7. Ratios were reported only when both means were positive. Complete eligible fixed-window pairs were available for 36 cultures, so all four endpoints were additionally compared within that same 36-culture subset. Calibration-dependent O_2_ sensitivity analyses excluded the three cultures with a pre-initial-illumination median ≥10 µM, and separately excluded any culture with such a flag before cycle 1 or cycles 5–7. The primary cohort was retained for the main figures. Paired resampling of cultures (Efron, 1979) provided descriptive empirical-bootstrap intervals for mean differences and ratios of means (Tables S10–S13).

## Results

### H_2_ responses across the irradiance series

Our irradiance-response dataset covered 54 biological replicates from 17 strains, with one experimental series per strain (Figure S5). The irradiance giving the highest mean peak H_2_ rate differed from strain to strain: seven strains peaked at 1110 µmol photons m⁻² s⁻¹, four at 722, three at 357, two at 107, and one at 52 µmol photons m⁻² s⁻¹. For two strains, G1098 and G1105, the highest mean response was less than 5% above the next-highest one. For 13 strains, the irradiance chosen for the repeated-cycle experiments matched this measured peak; for G1098 and G1105, we instead used the irradiances already available, as listed in Table S3.

### H_2_ responses during repeated light and dark cycles

The repeated-cycle dataset covered 48 cultures from 15 strains. The H_2_ traces varied both in overall response size and in how different the first exposure was from later ones (Figure 1). Using the early-response calculation shared by Figures 1 and 2, the first-light rate exceeded the mean of cycles 5–7 in 43 of the 48 cultures (Table S4). The ratio of each strain’s mean first rate to its mean later rate was above 1 in 14 of the 15 strains, ranging from 0.63 in G1122 to 17.48 in G1095.

The three largest ratios were in G1095 (17.48), G1107 (13.15) and G1096 (4.72). Their first and later rates (mean ± SD) were 10.39 ± 7.33 and 0.59 ± 0.26 µM min⁻¹ for G1095 (n = 3); 6.42 ± 2.48 and 0.49 ± 0.06 µM min⁻¹ for G1107 (n = 3); and 20.03 ± 7.86 and 4.25 ± 3.81 µM min⁻¹ for G1096 (n = 4). In all ten cultures across these three strains, the first rate was higher than the later rate. By contrast, G1093 had a higher mean first rate — 56.64 µM min⁻¹ — but a much smaller ratio of 1.67. So a large first-to-later ratio did not simply track a strain’s absolute initial H_2_ rate (Figure 2).

### Smaller contrasts and dependence on the rate summary

G1098 and G1104 had first-to-later ratios of 2.00 and 1.76. G1097 and G1103 were closer to 1 (1.12 and 1.01): their first rate was higher than the later mean in two of three cultures for G1097, and in one of two for G1103. G1122 went the other way — all three of its cultures had a higher later rate, with the mean rising from 8.80 ± 4.09 to 13.86 ± 2.06 µM min⁻¹. G1101 had a ratio of 1.36, but its rates varied a lot relative to their means (first 2.00 ± 2.34; later 1.47 ± 2.10 µM min⁻¹; n = 3). Table S4 gives the underlying paired values.

Higher first sustained-light rates also predominated under alternative endpoints, although the magnitude and some culture-level directions depended on the time window (Table S10). Using a common ratio-of-means summary, the full-period peak-30-s analysis gave higher first rates in 43 of 48 cultures and ratios above 1 in 14 of 15 strains, as did the primary early-response analysis. Whole-120-s slopes were complete for all 48 cultures and gave higher first rates in 46 cultures and ratios above 1 in 14 strains. G1122 retained a strain ratio below 1 (0.854), but only two of its three cultures retained the inverted direction. Fixed-first-30-s estimates passed the coverage checks in 36 cultures; all 36 had a higher first slope, but 22 had a nonpositive later slope, precluding a positive first-to-later ratio. G1122 therefore did not retain its inversion for this onset-defined endpoint. Comparing all four endpoints within the same 36 cultures confirmed that these differences were not solely due to missing fixed-window estimates.

Exploratory analysis of log culture-level H_2_ ratios detected strain-associated variation (one-way ANOVA, F(14, 33) = 9.53, p = 6.44 × 10⁻⁸). Strain accounted for most of this variance (η² = 0.80; intraclass correlation 0.73, or 0.53 after omitting the three strains with the largest ratios). The median-centered Brown–Forsythe variance check indicated unequal variances (p = 0.031), and Welch’s analysis retained the association (F(14, 11.60) = 15.33, p = 2.04 × 10⁻⁵). After omitting G1095, G1096 and G1107, the Welch result remained detectable (p = 0.00854). Games–Howell comparisons identified 16 of 105 pairs with adjusted p < 0.05 (Table S13). Every such pair involved G1095, G1107 or G1122; none involved G1096, whose culture-level ratios were variable, and none compared two of the remaining eleven strains. The current data support heterogeneous strain-associated responses rather than a fully resolved ranking.

### O_2_ responses during repeated illumination

The full-period cycle analysis showed O_2_ rates rising in several strains (Figure S9), alongside changes in the concentration profiles (Figure S3). In several strains, reconstructed bulk O_2_ sometimes exceeded 10 µM during or shortly after illumination and often fell below the operational threshold again within about 1–2 min of darkness. Recovery was not universal. In G1107 at 722 µmol photons m⁻² s⁻¹, the median rate rose from 0.13 µM min⁻¹ in cycle 1 to 7.37 µM min⁻¹ in cycle 7. In G1095 at 1110 µmol photons m⁻² s⁻¹, it rose from 2.36 in cycle 1 to 10.45 in cycle 5, then was 6.80 µM min⁻¹ in cycle 7. In G1093 and G1097, the reconstructed pre-light O_2_ median was above our 10-µM threshold before 16 later illuminations, including seven within cycles 5–7; three initial medians were also above it, even though our anoxia check at the time had judged them anoxic (Table S5). We kept these calibration-dependent discrepancies as they were, rather than forcing the baselines to zero. This means the later light exposures carry different O_2_ histories, and not all of them can be treated as an equivalent return to anoxia.

Removing the three cultures with initial pre-illumination O_2_ medians ≥10 µM left higher first rates in 40 of 45 cultures and ratios above 1 in 13 of 15 strains (Table S12). The G1097 ratio changed from 1.117 to 0.930; the primary high-response strains and the inverted primary response in G1122 were unaffected. Requiring every retained culture to have unflagged starts in cycle 1 and cycles 5–7 left 44 cultures, with higher first rates in 39 cultures and ratios above 1 in 13 strains. This stricter analysis left only one culture each for G1093 and G1097, so their sensitivity estimates are descriptive.

### CO_2_ responses during repeated illumination

CO_2_ responses also differed between the first illumination and the last three cycles (Figure S4). On the nominal conversion scale described in the Methods, using the early-response 30-s regressions shown in Figure S4, the group-mean light-period slope was negative in 13 of the 15 strains during the first exposure, and in all 15 strains by cycles 5–7. Two strains, G1093 and G1094, instead had a positive mean first-exposure slope (0.081 and 0.065 nominal units min⁻¹) followed by a negative late-cycle mean (−0.635 and −0.247 nominal units min⁻¹); because the sign flipped, we did not report a first-to-later CO_2_ ratio for these two. Among the remaining strains, the ratio of the mean first-exposure slope to the mean cycles-5–7 slope ranged from 0.14 in G1101 to 2.10 in G1098. G1107 showed a larger negative slope later on (−0.287 vs. −0.054 nominal units min⁻¹ initially), whereas G1095’s first and later values were similar (−0.325 and −0.282 nominal units min⁻¹). Figure S10 shows full-period rate estimates across all seven cycles separately.

## Discussion

### A widespread but variable first-light response

An enhanced first-light H_2_ response, previously characterized in *C. reinhardtii* (Milrad et al., 2018, 2023), was the predominant pattern across this collection. First sustained-light rates exceeded later rates in 14 of 15 strains, representing the *Chlorellaceae*, *Scenedesmaceae*, *Selenastraceae* and *Chlamydomonadales*, and this predominance held across alternative rate windows and O_2_-based exclusions. Its magnitude varied widely, from near unity in G1097 and G1103 to more than 13 in G1107 and G1095, and strain accounted for most of the variance in culture-level ratios. G1122 was the exception, with higher later rates in all three cultures; this reversal was retained by the whole-period strain mean but not by a fixed-onset 30-s endpoint, and the near-unity G1097 contrast was sensitive to O_2_-based exclusion. Because pairwise comparisons resolved only the extreme strains, the intermediate strains should not be ranked. Figure 2 distinguishes the first-to-later contrast from the initial net H_2_ rate. G1095 had the largest ratio because its later rate was small relative to its first rate, whereas G1093 and G1102 had larger initial rates. The ratio describes the change within the recorded sequence; it does not measure cumulative H_2_ yield, hydrogenase abundance or an intrinsic species-level production capacity. Paired culture values in Table S4 show the consistency and spread of these responses at the available sample sizes.

Recorded provenance provides context for ecological hypotheses (Table S9). G1095 and G1107, with the largest primary H_2_ ratios, originated from the Yarkon River and a winter puddle near Bental reservoir, respectively; G1103, with a primary ratio near unity, came from the El-Al Stream. Other Yarkon isolates, G1094 and G1104, had ratios of approximately 1.99 and 1.76. Locality therefore does not consistently predict the contrast within this collection. Dissolved O_2_ at collection was not measured, so adaptation to habitat anoxia could not be tested.

### Possible physiological explanations

Studies in *C. reinhardtii* suggest several possible explanations. Competition for electrons can terminate H_2_ production while hydrogenase remains active (Milrad et al., 2018). Hydrogenase activity and cyclic electron flow contribute to carbon-fixation induction (Godaux et al., 2015), while O_2_ photoreduction also participates in this transition (Burlacot et al., 2018). Bidirectional H_2_-linked electron transfer can affect responses to changing light (Milrad et al., 2021). Differences in the timing or balance of these processes could contribute to strain variation. Paired changes in H_2_, O_2_ and CO_2_ provide a basis for testing these hypotheses in individual strains.

Milrad et al. (2023) linked attenuation after illumination to PSII photosynthetic control and observed recovery after 15 min of dark anoxia. G1098 is the same *C. reinhardtii* CC124 reference strain. Its primary ratio here was 2.00, in the same direction as the earlier approximately threefold contrast. Differences in irradiance (722 µmol photons m⁻² s⁻¹ here versus 370 previously), cycle number, fitting and averaging definitions, and growth conditions may account for the smaller contrast. Because the 3-min dark intervals used here are shorter than this recovery time, later cycles likely reflect incomplete recovery, consistent with the continued decline of some later rates between cycles 5 and 7 (Supplementary Results). Whether the same PSII mechanism operates in the other strains can be tested by varying the dark interval between illuminations and by applying the spectroscopic and biochemical approaches used for *C. reinhardtii*.

The measurement sequence combined an initial 1-h dark incubation with 3-min dark intervals between sustained-light periods. Dark adaptation, recent photosynthetic activity, metabolite pools and cumulative O_2_ exposure therefore changed together. A larger first response could reflect a state established during prolonged darkness, incomplete recovery between short cycles, regulation triggered by the preceding illumination, or a combination of these effects.

### What the accompanying gases show

HydA1 is highly O_2_-sensitive in cell-free assays: recombinant *C. reinhardtii* HydA1 in solubilized *E. coli* samples showed 50% activity loss in less than 1 s of exposure to atmospheric O_2_ (21%; Cohen et al., 2005). However, intact cells can maintain H_2_ production at much higher bulk O_2_ through local O_2_ scavenging (Liran et al., 2016). The 10 µM criterion used here was therefore a pragmatic operational cutoff, not a measured species-specific inactivation threshold. We considered brief excursions followed by recovery below 10 µM during darkness acceptable for retaining the affected cycles, while recognizing that recovery of bulk O_2_ does not establish that hydrogenase activity was unaffected. Several strains, most clearly G1107 and G1095, showed greater net O_2_ accumulation in later cycles while H_2_ rates declined. Net O_2_ change combines production and consumption and cannot be equated with gross PSII activity. Three reconstructed initial baselines and some later pre-illumination medians exceeded the operational threshold (Table S5), so not all recordings started from the same O_2_ conditions; the predominance of higher first rates was nonetheless retained when the flagged cultures were excluded. No cross-strain association was detected between H_2_ and O_2_ first-to-later ratios (Spearman ρ = 0.007, n = 15, Holm-adjusted p = 0.985; Table S8). This comparison concerns net O_2_ exchange and does not resolve the mechanistic contribution of O_2_ to the H_2_ response. CO_2_ was recorded on a nominal signal scale; quantifying inorganic-carbon flux would additionally require treatment of CO_2_–bicarbonate interconversion (Douchi et al., 2019). Within these limits, the positive first slopes followed by negative later slopes in G1093 and G1094 are consistent with a delayed onset of carbon assimilation during the first illumination. Across the 13 strains with negative first and later mean CO_2_ slopes, H_2_ and CO_2_ ratios were not significantly correlated (ρ = 0.363, Holm-adjusted p = 0.447; Table S8).

### Scope and analytical limitations

These comparisons apply to the strains and conditions tested. Irradiances differed among strains, while increasing irradiance in the sequential series was coupled to elapsed time and prior light exposure. In G1098 and G1105, the repeated-cycle irradiance did not match the sampled irradiance-response maximum. Instrument, acquisition date and strain were not fully separated by the design. Four strain identities remain based on morphology, and several other assignments are provisional. The observed range therefore cannot be interpreted as an intrinsic species ranking, a phylogenetic pattern or a response independent of irradiance.

Rate selection emphasizes different parts of the transient. The fixed-onset, whole-period and peak-window comparisons use the same ratio-of-means summary and a common-culture subset, making endpoint dependence explicit. Late weak fits can remain uncertain even with reproducible window selection; 15 of 144 primary late H_2_ fits had R² below 0.5. Culture resampling illustrates this uncertainty — for example, G1096’s ratio of 4.72 had an empirical-bootstrap range of 1.94–16.72 (Table S11) — but cannot correct for the sparse n = 2–5 support, physical gas loss or absolute-calibration error. Intermediate strain differences and mechanistic contributions remain unresolved within this study’s descriptive scope.

Finally, MIMS records the net change in dissolved gas while itself withdrawing gas through its inlet membrane. Without an independent, gas-specific correction for that withdrawal, we cannot quantitatively separate true physical loss from simultaneous biological production and consumption. A constant calibration offset does not affect a slope, and a stable, multiplicative sensitivity factor cancels out of a first-to-later ratio within one recording — but a sensitivity that drifts over time, or a loss that depends on concentration, does not necessarily cancel. For these reasons, we favor interpreting the reproducible patterns and the paired absolute rates alongside the ratios, rather than treating the ratios as exact measurements of a regulatory mechanism.

## Conclusions

The first-light enhancement of H_2_ production described in *C. reinhardtii* was the predominant response across 15 green microalgal strains from four lineages, and it persisted under the primary early-response endpoint and several alternative summaries. Its magnitude, however, varied by more than an order of magnitude among strains. Exact magnitudes, G1122’s reversal and near-unity contrasts depended on the window or O_2_-based cohort definition. O_2_ and nominal CO_2_ records added physiological context. Strains with exceptionally strong (G1095, G1107) or reversed (G1122) responses are natural candidates for testing whether PSII photosynthetic control explains this variation.

## Acknowledgments

We thank Eyal Zaksh for his contribution to this work and for assistance with obtaining algal samples.

## Funding

This study was supported by the Israel Science Foundation (grant 941/22), the Ministry of Science and Technology (grant 8395), and the Council for Higher Education (VATAT) Excellence Center for Energy.

## Conflicts of Interest

The authors declare no conflict of interest.

## Data and code availability

The numerical data underlying the figures, tables and statistical analyses are provided in Supplementary Data 1 as an Excel workbook. Supplementary Data 2 contains the 102 raw MIMS ASCII recordings, culture-to-file manifest, calibration coefficients, analysis settings, calibrated trace and rate exports, statistical scripts and figure files. The archive includes 48 repeated-cycle and 54 irradiance-response cultures, with SHA-256 checksums linking each raw recording to its original export. These files support reproduction of the figures and sensitivity analyses.

## AI assistance

OpenAI Codex and Claude (Anthropic) were used for language editing, analysis of the recordings, figure formatting and document preparation. The numerical data are included in Supplementary Data 1; the analysis scripts, source mappings and figure-generation files are included in Supplementary Data 2.

## Supplementary information

**Table S1:** Working taxonomic assignments and identification evidence for the 17 strains. Molecular support is not treated as definitive species identity. G1099, G1100, G1106 and G1122 retain morphology-only assignments. The companion reanalysis adds the qualifications below; provenance for the full MIMS collection is given in Table S9. G1106 and G1121 contribute irradiance-response data only.

| Code | Identification | Identification basis | Dataset |
| --- | --- | --- | --- |
| G1093 | *Chlorella ohadii* | Reference laboratory culture linked by provenance to SAG 2657; rDNA does not distinguish *C. ohadii* independently | Switch and PI |
| G1094 | *Chlorella sorokiniana* | Morphology and rDNA (previous study) | Switch and PI |
| G1095 | *Micractinium tetrahymenae* | Morphology and rDNA; provisional species assignment in companion reanalysis | Switch and PI |
| G1096 | *Desmodesmus abundans* | Morphology and rDNA (previous study) | Switch and PI |
| G1097 | *Tetradismus obliquus* | Morphology and rDNA (previous study) | Switch and PI |
| G1098 | *Chlamydomonas reinhardtii* | Reference strain CC124 | Switch and PI |
| G1099 | *Chlorella* sp. | Morphology only | Switch and PI |
| G1100 | *Chlorella sorokiniana* | Morphology only | Switch and PI |
| G1101 | *Chlorolobion braunii* | Morphology and rDNA; provisional species assignment in companion reanalysis | Switch and PI |
| G1102 | *Acutodesmus obliquus* | Supplied as SAG 276-3d; *Acutodesmus obliquus* is synonymous with *Tetradismus obliquus* | Switch and PI |
| G1103 | *Lobochlamys segnis* | Morphology and rDNA (previous study) | Switch and PI |
| G1104 | *Monoraphidium cf. dybowskii* | Morphology and rDNA; tentative cf. assignment | Switch and PI |
| G1105 | *Micractinium* sp. | Morphology and rDNA; generic placement unresolved | Switch and PI |
| G1106 | *Stichococcus bacillaris* | Morphology only | PI only |
| G1107 | *Desmodesmus abundans* | Morphology and rDNA (previous study) | Switch and PI |
| G1121 | *Chlorella lewinii* | Morphology and rDNA; provisional species assignment in companion reanalysis | PI only |
| G1122 | *Monoraphidium convolutum* | Morphology only | Switch and PI |

“sp.” denotes a genus-level working name and “cf.” a tentative species assignment; unresolved generic placements remain qualified. G1097 and G1102 are distinct cultures despite synonymous supplied species names. Collection cultures were selected visually as unialgal; axenicity was not tested. Taxonomic and provenance information follows the companion characterization manuscript.

**Table S2:** Recorded affine coefficients for raw ion-current conversion, C = mI + b for H_2_/O_2_ and N = mI + b for nominal CO_2_ signal N. I is in A. H_2_/O_2_ scale units are µM, with slopes µM A⁻¹; CO_2_ scale units and slopes are nominal units and nominal units A⁻¹. Coefficients do not independently validate absolute accuracy. Table S3 maps profiles to the retained experiments. C1 is a switch profile from one MIMS system; C2 is a later profile from that same system, with the recorded H_2_ and O_2_ background offsets subtracted once. C3 is the profile from the second MIMS system. C4a and C4b are irradiance-response profiles that share the same H_2_/O_2_ coefficients and CO_2_ slopes but have different CO_2_ intercepts. Table S3 shows which profile applies to each selected experiment. We kept these coefficients exactly as recorded during acquisition. Separate profiles were needed because we used two instruments and replaced filaments during the study. We applied no additional offset and no argon factor. A constant concentration offset does not change the slope of a concentration trace, but absolute concentrations and O_2_-threshold calls depend on the calibration intercept as well as the slope.

| Profile | Gas | m (scale units $A^{-1}$) | b (scale units) |
| --- | --- | --- | --- |
| C1 | $H_2$ | 8.569748917e+10 | 0.8569748917 |
| C1 | $O_2$ | 5.800390731e+10 | -2.675656263 |
| C1 | $CO_2$ | 1.141821118e+11 | -17.72974780 |
| C2 | $H_2$ | 7.222222222e+10 | -12.78333333 |
| C2 | $O_2$ | 8e+10 | -13.12 |
| C2 | $CO_2$ | 3.528365387e+10 | 0 |
| C3 | $H_2$ | 9.193636533e+11 | -132.116004 |
| C3 | $O_2$ | 6.281224011e+12 | -12.13035397 |
| C3 | $CO_2$ | 1e+11 | 0 |
| C4a, C4b | $H_2$ | 8.09e+10 | 68.83256524 |
| C4a, C4b | $O_2$ | 5.800390731e+10 | -2.675656263 |
| C4a | $CO_2$ | 3.528365387e+10 | -83.81209967 |
| C4b | $CO_2$ | 3.528365387e+10 | 0 |

**Figure S1:**
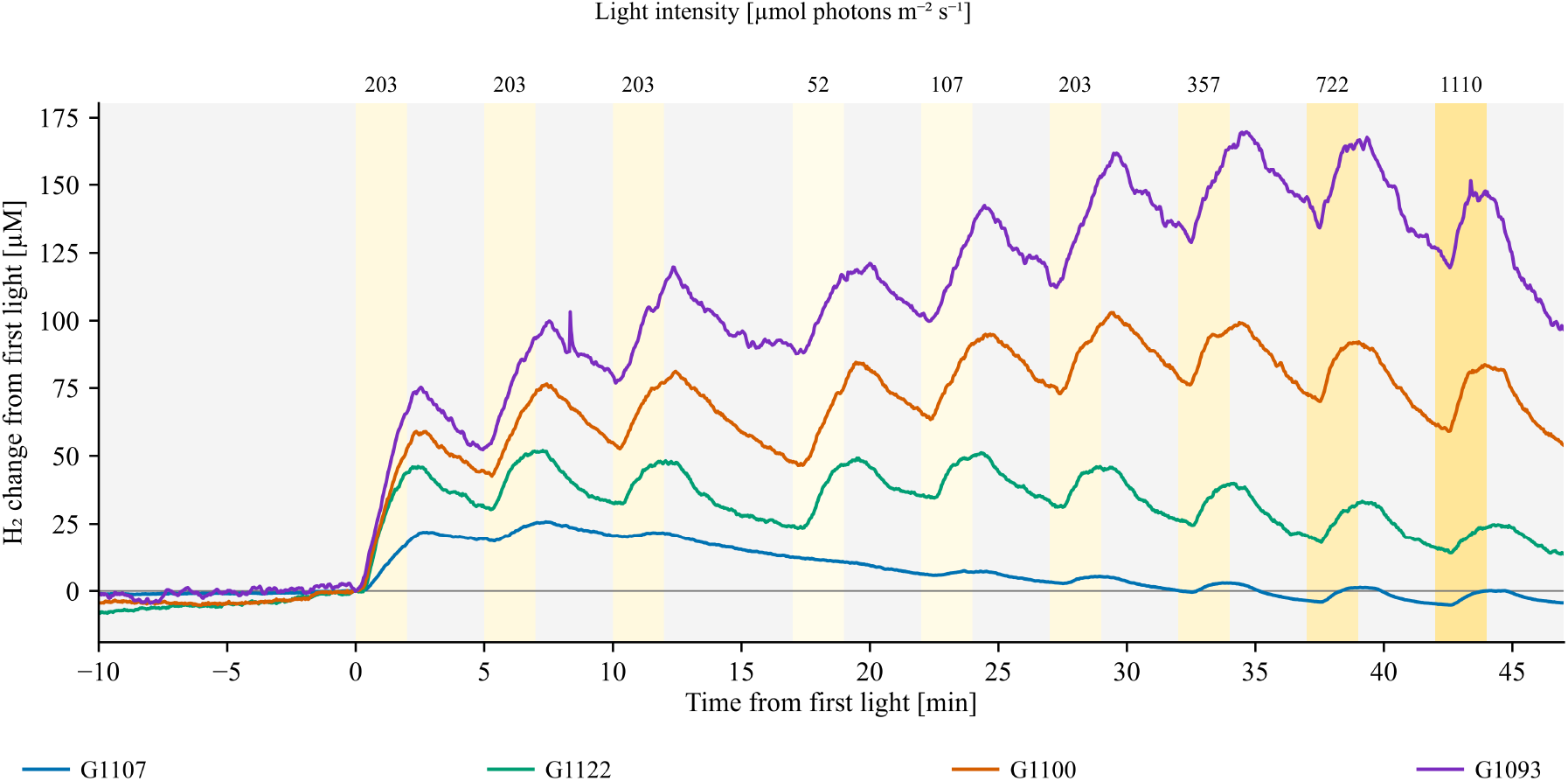
Example H_2_ recordings illustrating the sequential irradiance-response protocol. Each colored line is one individual culture of G1107, G1122, G1100 or G1093. We show the complete trace relative to its interpolated H_2_ concentration at first light onset, without rescaling its amplitude or aligning individual cycles. Three 2-min conditioning light periods at 203 µmol photons m⁻² s⁻¹ came first; after the final 5-min dark interval, six 2-min light periods followed at 52, 107, 203, 357, 722 and 1110 µmol photons m⁻² s⁻¹. Irradiances are shown above the light intervals; yellow shading gets stronger with irradiance, and gray marks darkness. These four examples illustrate the protocol; the complete 17-strain rate summaries are shown in Figures S5–S7. First exposure denotes the first sustained 2-min measurement light period in the analyzed sequence.

**Figure S2:**
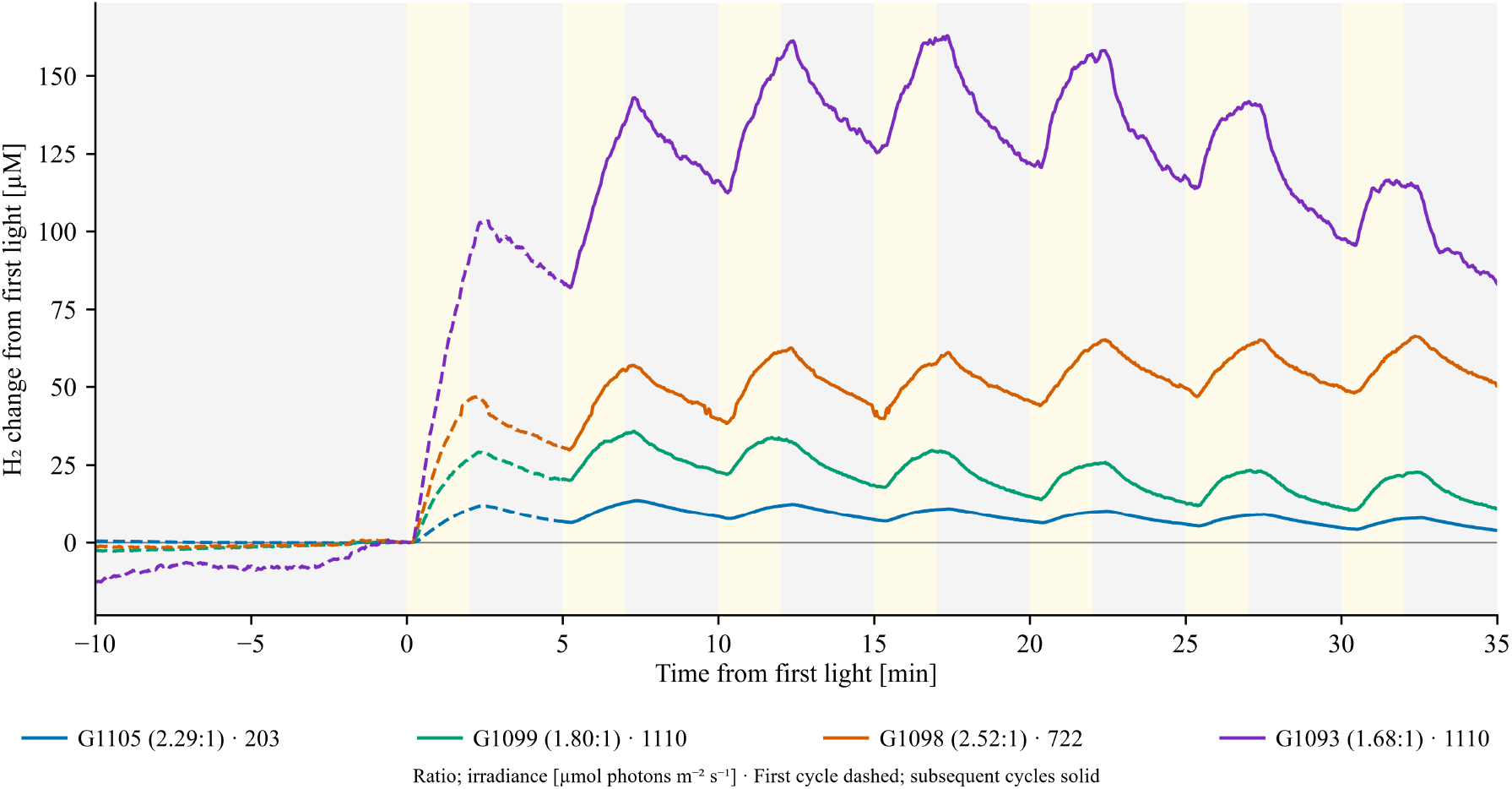
Example H_2_ recordings during seven repeated light–dark cycles. Each colored line is one individual culture of G1105, G1099, G1098 or G1093, at 203, 1110, 722 or 1110 µmol photons m⁻² s⁻¹ respectively. We show H_2_ changes relative to the concentration at first light onset, without aligning cycle by cycle. The dashed segments cover the pre-light baseline and first light–dark cycle; later cycles are solid. Yellow and gray mark the 2-min light and 3-min dark intervals. The ratios in the line key are the first-light rate divided by the mean light rate in cycles 5–7 for that individual recording, using the early-extremum 30-s method. These four examples illustrate the full protocol; they are not replicate averages. The complete 15-strain cycle-rate summaries are shown in Figures S8–S10. First exposure denotes the first sustained 2-min measurement light period in the analyzed sequence.

**Figure S3:**
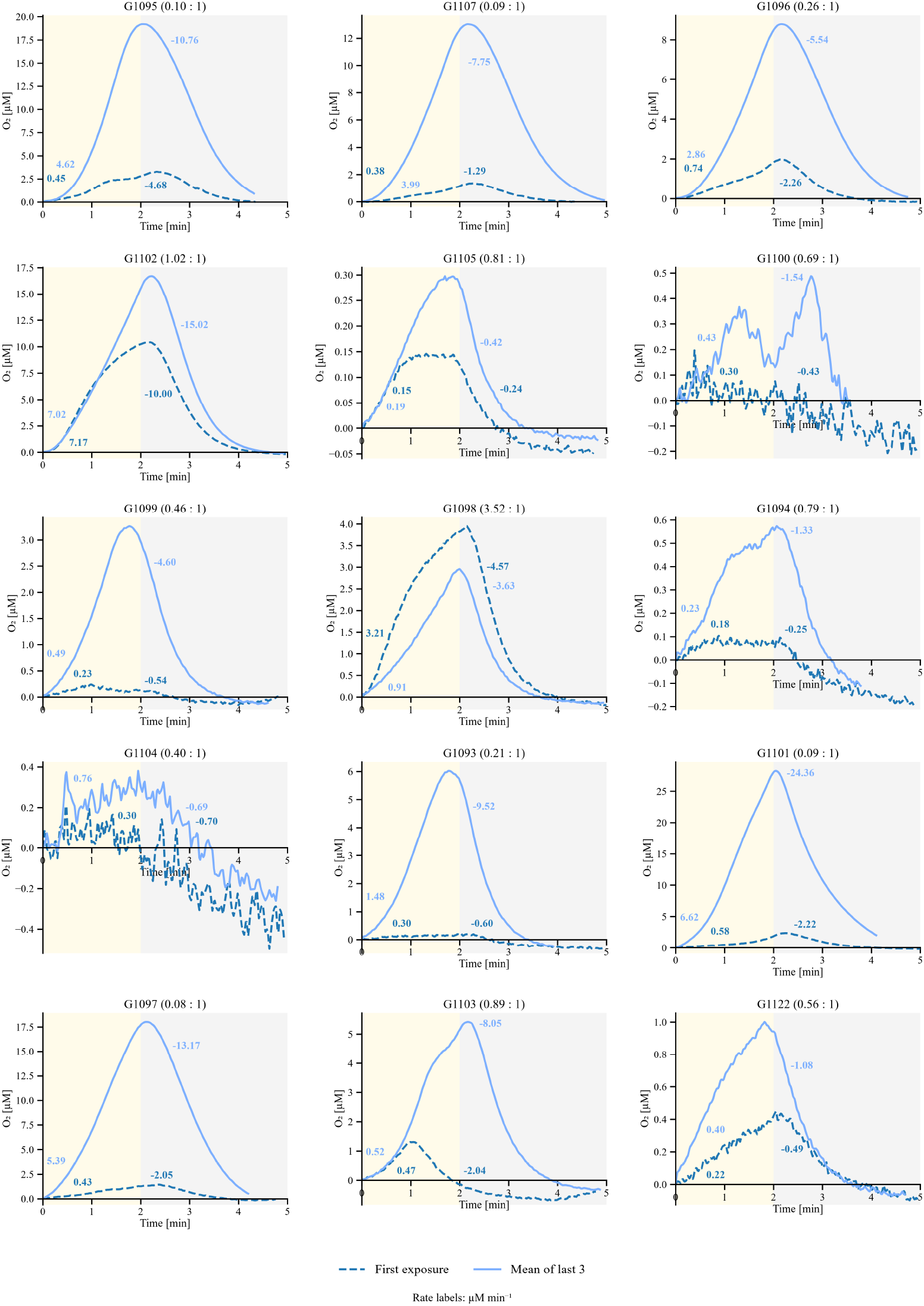
First and later O_2_ responses during repeated illumination. Panels show the same 15 strains and 48 cultures as Figure 1, in the same H_2_-based order (irradiances and n, Table S3). The dark-blue dashed line is the mean first exposure, and the light-blue solid line is the mean of exposures 5–7, averaged first within cultures and then across cultures. We shifted each exposure to a common time-and-concentration origin, so the O_2_ values shown are changes from that origin, not evidence of absolute anoxia. Yellow and gray mark the nominal light/dark schedule after alignment, not the exact lamp transitions. The four numbers are mean early-extremum 30-s rates in light and darkness (µM min⁻¹); the panel ratio compares the mean first and later light rates. No error bands are shown here; Figure S9 gives the full-phase 30-s cycle-rate summaries. We show a panel ratio only when both mean light rates are positive. First exposure denotes the first sustained 2-min measurement light period in the analyzed sequence.

**Figure S4:**
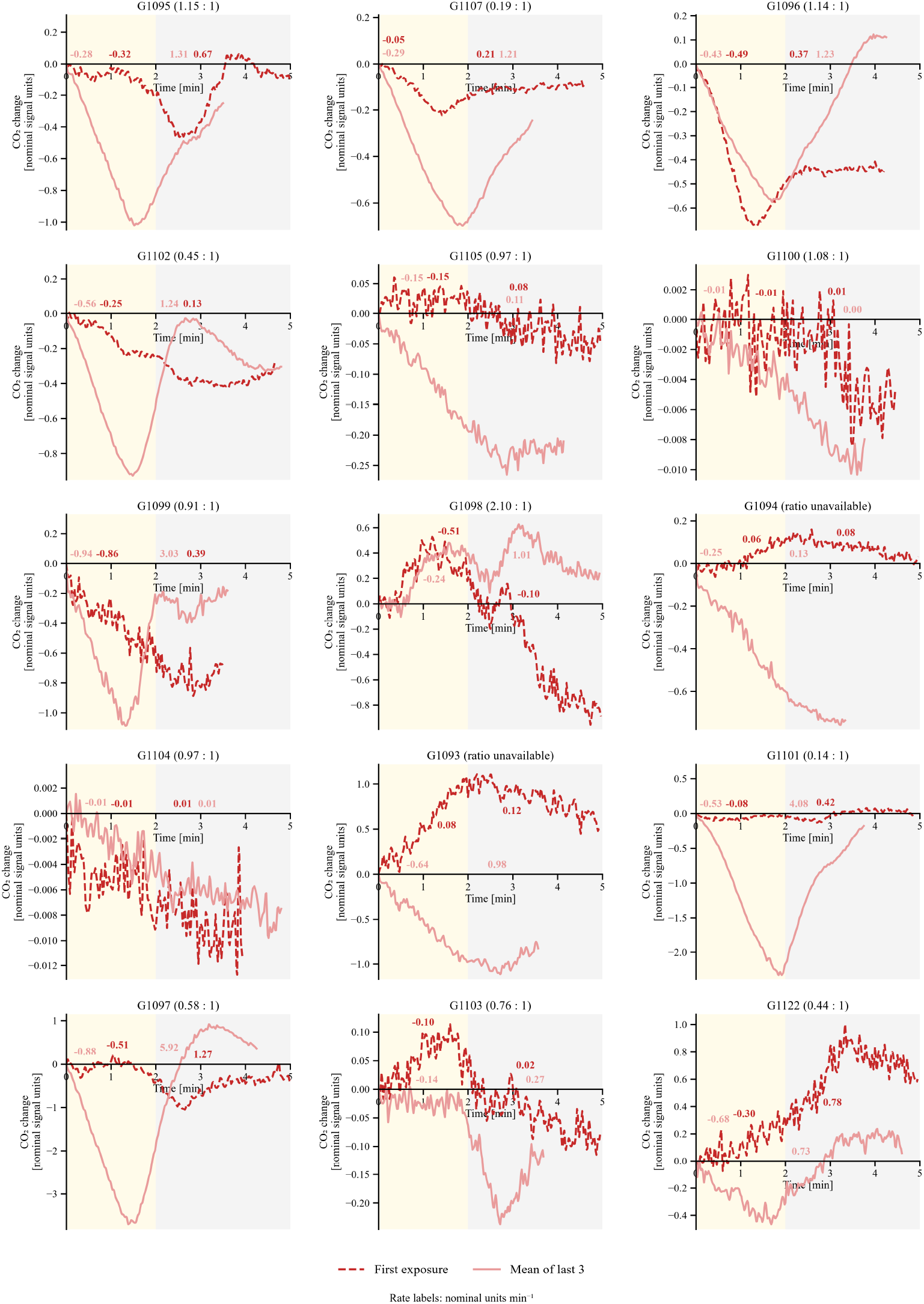
First and later CO_2_ responses during repeated illumination. Panels show the same 15 strains and 48 cultures as Figures 1 and S3, ordered by the H_2_ ratio (irradiances and n, Table S3). The dark-red dashed line is the mean first exposure, and the light-red solid line is the mean of exposures 5–7, with each culture weighted equally. We shifted each exposure to its detected response maximum before a decline (or to a documented default onset when no maximum could be resolved) and to the matching concentration baseline. The values show change in dissolved CO_2_, with negative rates meaning a net decline and positive rates a net increase. Yellow and gray mark the nominal light/dark schedule after alignment, not the exact lamp transition of each recording. The numbers give mean early-extremum 30-s rates (nominal units min⁻¹). Where both exist, the panel ratio divides the negative mean first-light rate by the negative mean later-light rate, comparing the size of the net decline. Curves show no error bands. Figure S10 gives the full-phase cycle-rate summaries. CO_2_ axes use nominal signal units and nominal units min⁻¹ from the recorded coefficients; absolute concentrations and fluxes were not independently validated. First exposure denotes the first sustained 2-min measurement light period in the analyzed sequence.

**Table S3:** Peak irradiance and the irradiance selected for the repeated-cycle experiment, for each strain. “Peak irradiance” is the tested light level giving the highest mean 30-s H_2_ light-period rate in the selected irradiance-response series. Irradiance is given in µmol photons m⁻² s⁻¹. n is the number of biological replicates; Table S2 defines the calibration profiles. We kept the available switch experiments for G1098 and G1105 even though they do not match the measured PI peak. For G1098, the responses at 722 and 1110 µmol photons m⁻² s⁻¹ were close to each other (Figure S5). For G1097, we chose the series peaking at 722 µmol photons m⁻² s⁻¹, matched to its corresponding switch experiment. G1106 and G1121 contribute irradiance-response data only. PI stands for irradiance-response measurements. A dash means no switch recording is included. Different light–dark cycles from the same culture are repeated observations within one biological replicate, not separate replicates.

| Strain | Peak irradiance | Switch irradiance | PI n | Switch n | PI calibration | Switch calibration |
| --- | --- | --- | --- | --- | --- | --- |
| G1093 | 1110 | 1110 | 4 | 3 | C4a | C1 |
| G1094 | 722 | 722 | 3 | 3 | C4a | C2 |
| G1095 | 1110 | 1110 | 3 | 3 | C4a | C2 |
| G1096 | 1110 | 1110 | 3 | 4 | C2 | C2 |
| G1097 | 722 | 722 | 3 | 3 | C4a | C1 |
| G1098 | 1110 | 722 | 3 | 3 | C4a | C1 |
| G1099 | 1110 | 1110 | 3 | 3 | C4a | C1 |
| G1100 | 1110 | 1110 | 3 | 5 | C4b | C3 |
| G1101 | 1110 | 1110 | 3 | 3 | C4b | C2 |
| G1102 | 357 | 357 | 3 | 4 | C2 | C2 |
| G1103 | 722 | 722 | 3 | 2 | C4a | C2 |
| G1104 | 107 | 107 | 3 | 3 | C4a | C3 |
| G1105 | 107 | 203 | 3 | 3 | C4a | C1 |
| G1106 | 357 | - | 4 | - | C4b | - |
| G1107 | 722 | 722 | 4 | 3 | C2 | C2 |
| G1121 | 357 | - | 3 | - | C3 | - |
| G1122 | 52 | 52 | 3 | 3 | C4b | C1 |

**Figure S5:**
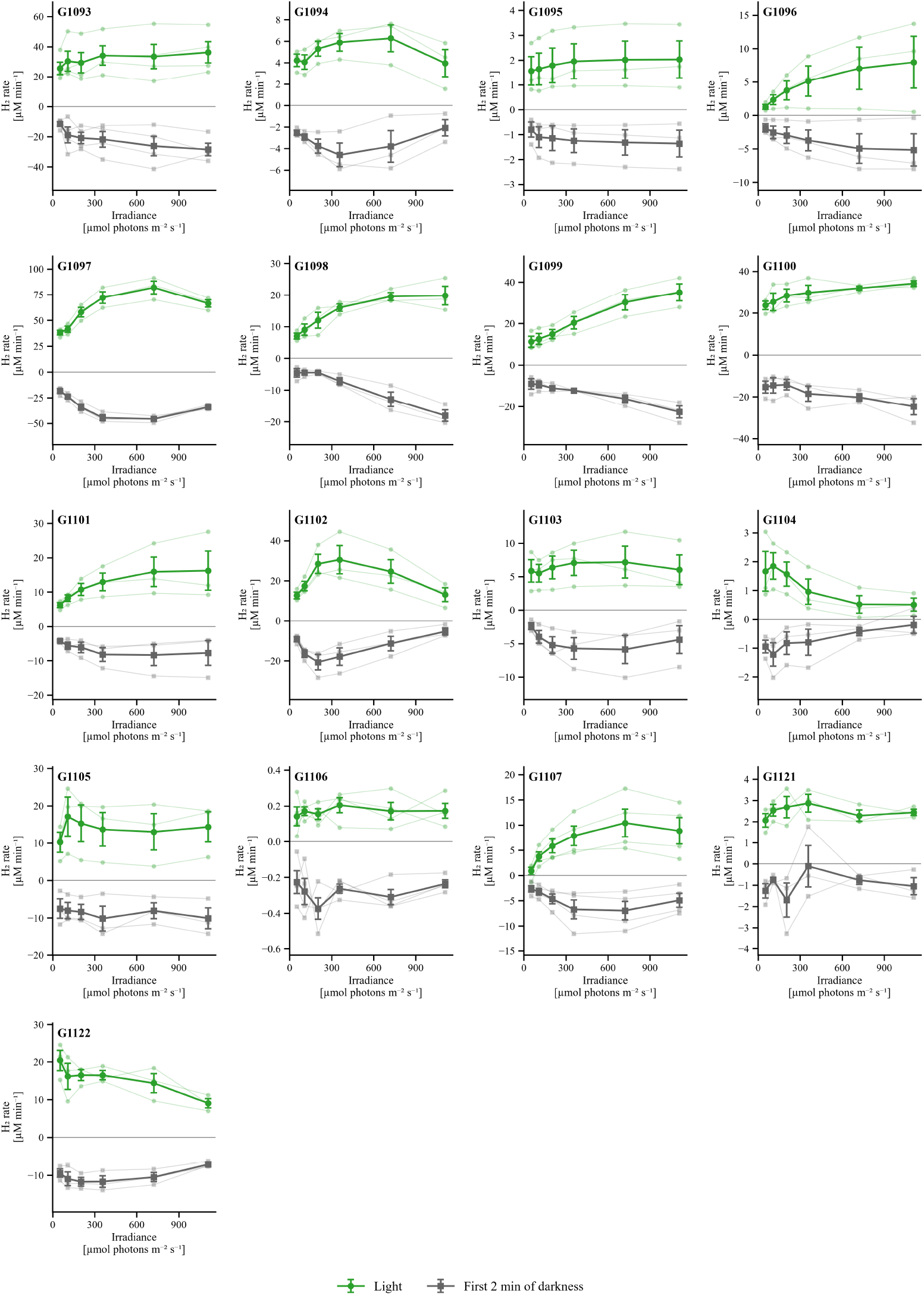
H_2_ rates across the sequential irradiance series. Panels show 17 strains and 54 independently grown cultures. Green circles are rates in light, and gray squares are rates during the first 2 min of the following dark interval. Thin pale lines connect individual-culture values; the prominent lines and error bars show arithmetic means ± SEM. For each phase, we selected the expected-sign 30-s slope from the full 2-min search described in the Methods; rates keep their signs. We applied the six irradiances in increasing order after three conditioning periods, which are omitted here. Positive values mean a net increase and negative values a net decrease in dissolved H_2_. Biological n = 4 for G1093, G1106 and G1107, and n = 3 for the other strains (Table S3). Panels follow numeric G-code order.

**Figure S6:**
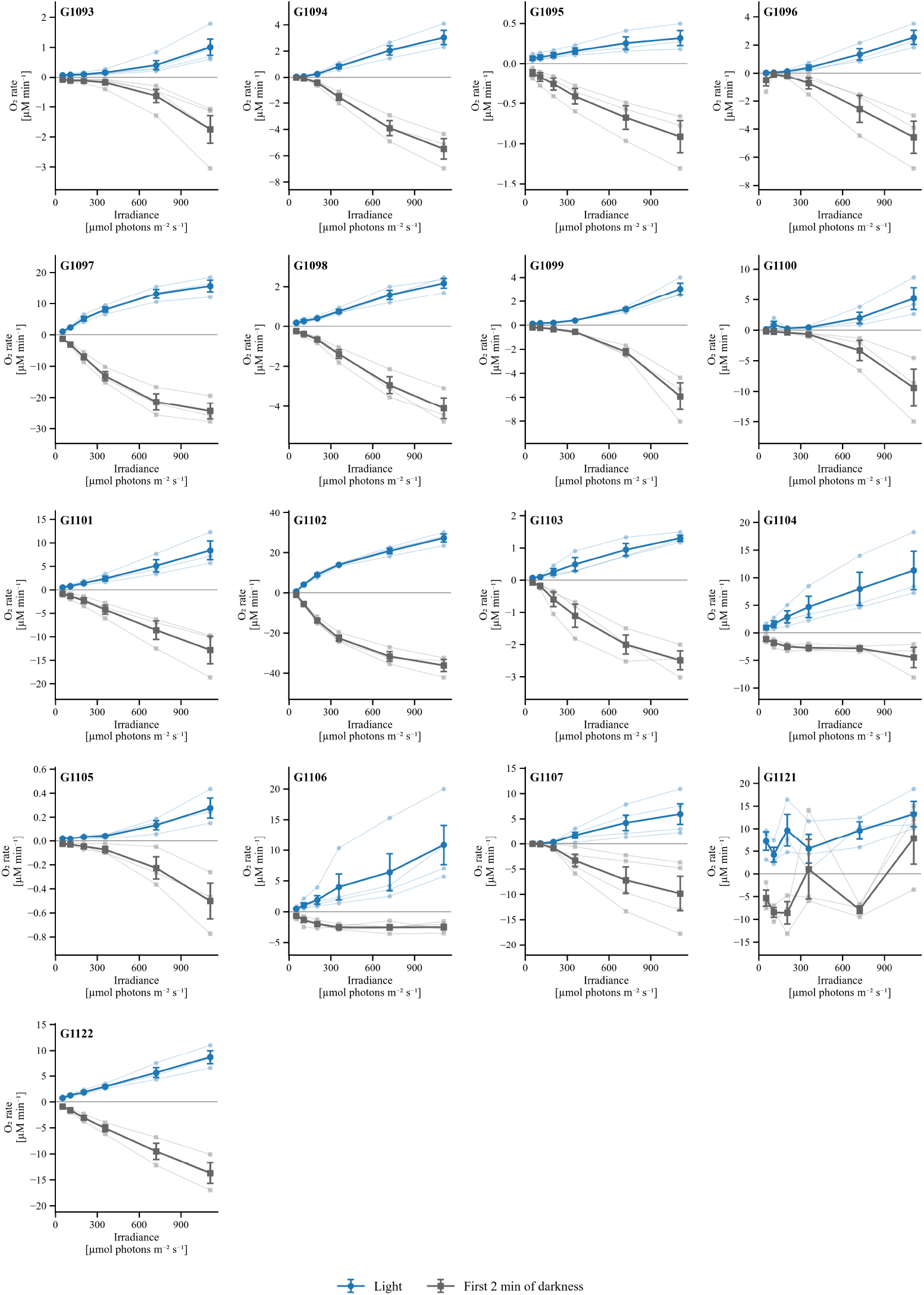
O_2_ rates across the sequential irradiance series. Panels show 17 strains and 54 independently grown cultures. Blue circles are light rates, and gray squares are rates during the first 2 min of the following dark interval. Thin pale lines connect individual cultures; the prominent lines and error bars show arithmetic means ± SEM. We selected rates from 30-s windows across each full 2-min analysis period, using the expected-sign rule described in the Methods. Positive and negative values mean net increases and decreases in dissolved O_2_. Only the six increasing test irradiances are shown; the conditioning periods are excluded. Biological n = 4 for G1093, G1106 and G1107, and n = 3 for the other strains (Table S3). Panels follow numeric G-code order.

**Figure S7:**
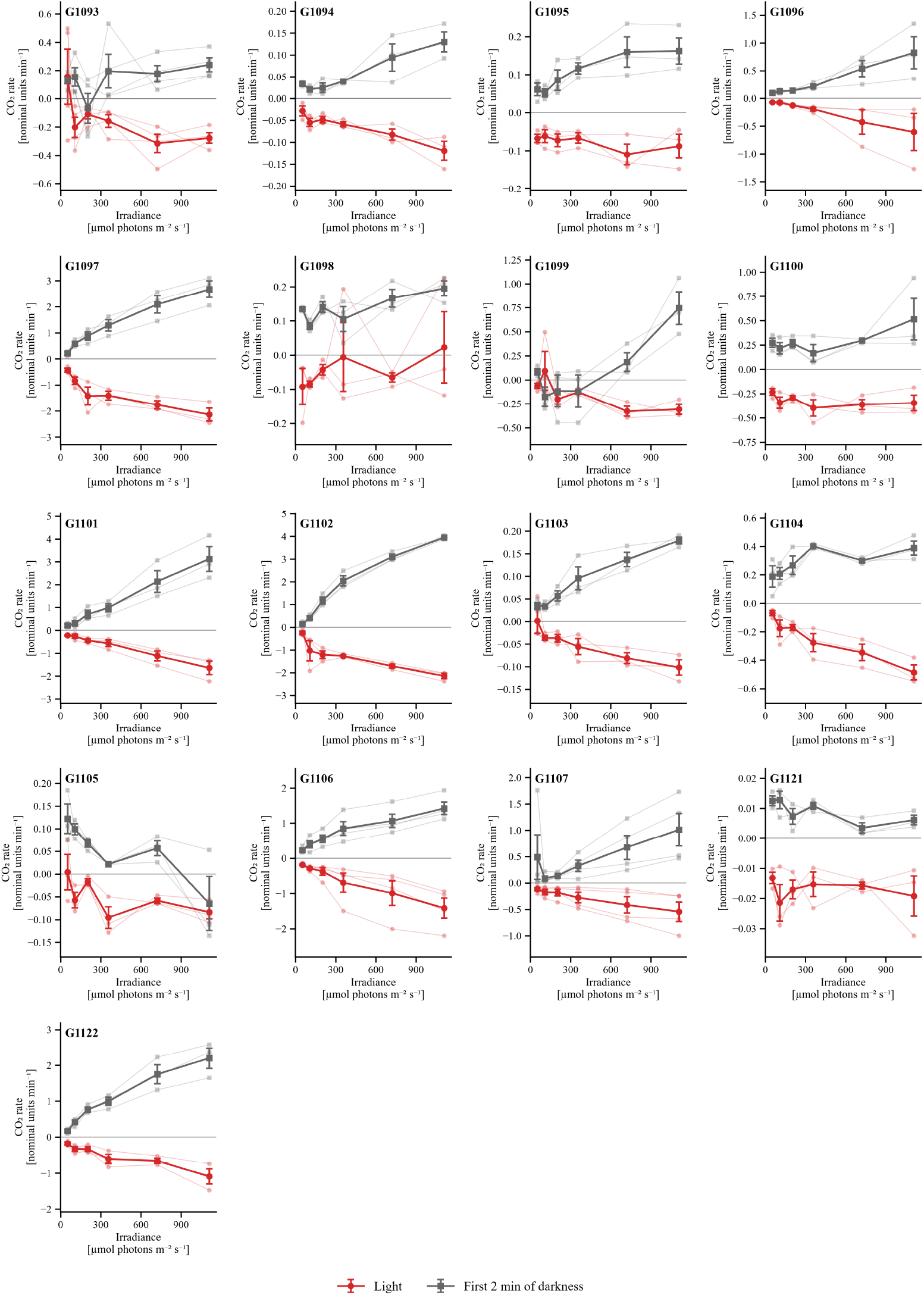
CO_2_ rates across the sequential irradiance series. Panels show 17 strains and 54 independently grown cultures. Red circles are light rates, and gray squares are rates during the first 2 min of the following dark interval. Thin pale lines connect individual cultures; the prominent lines and error bars show arithmetic means ± SEM. We obtained 30-s rates across each full 2-min analysis period, giving priority to negative slopes in light and positive slopes in darkness, as described in the Methods. Negative and positive values mean net decreases and increases in the CO_2_ signal. Only the six increasing test irradiances are shown; the conditioning periods are excluded. Biological n = 4 for G1093, G1106 and G1107, and n = 3 for the other strains (Table S3). Panels follow numeric G-code order. CO_2_ axes use nominal signal units and nominal units min⁻¹ on the recorded conversion scale, not independently validated absolute concentrations or fluxes.

**Figure S8:**
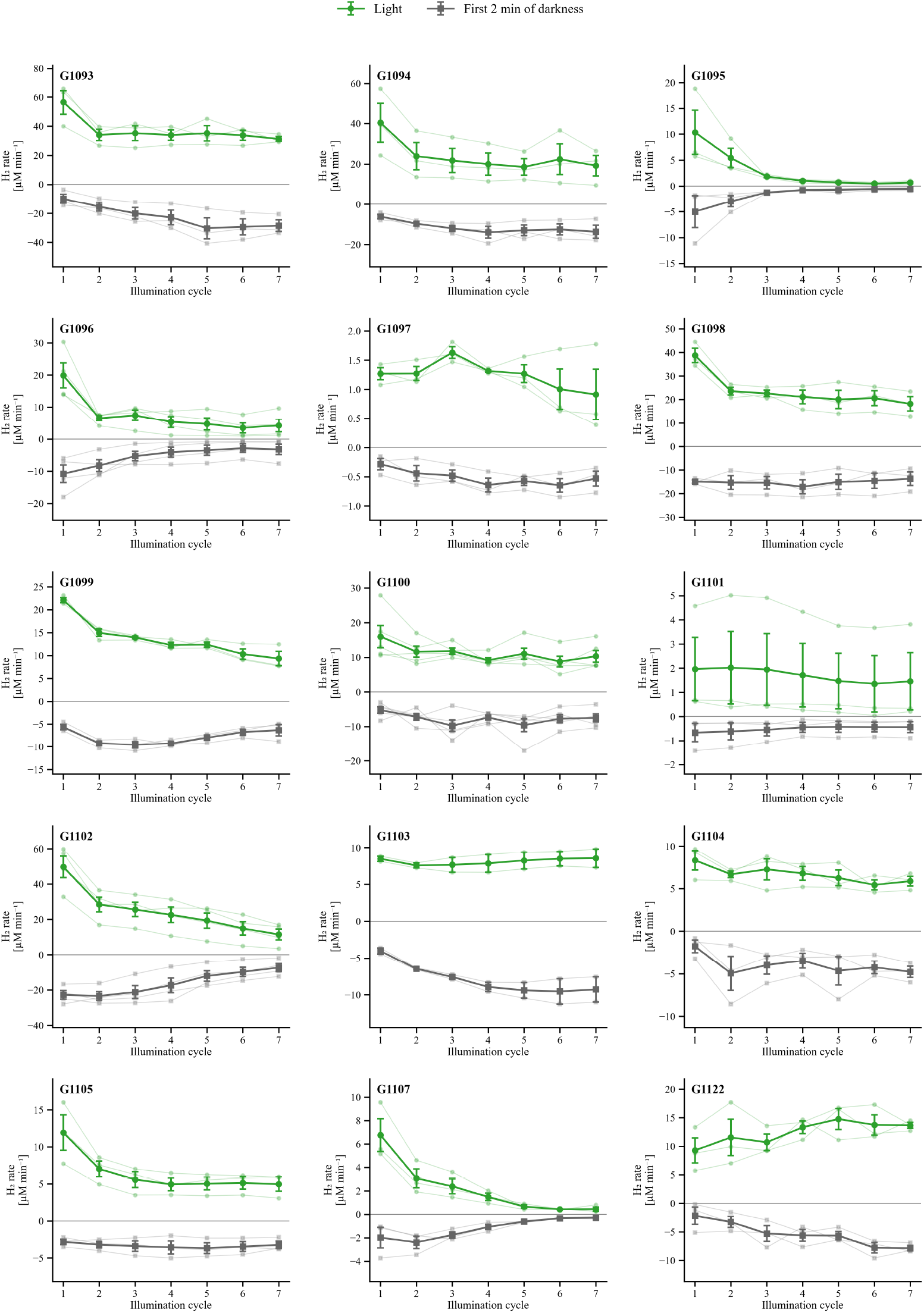
H_2_ rates through seven repeated light–dark cycles. Panels show 15 strains and 48 independently grown cultures, at each strain’s own constant irradiance (Table S3). Green circles are light rates, and gray squares are rates during the first 2 min of the following dark interval. Thin pale lines connect values from individual cultures; the prominent lines and error bars show arithmetic means ± SEM. We selected rates from 30-s windows across each full 2-min analysis period, using the signed selection rule described in the Methods. Positive and negative values mean net increases and decreases in dissolved H_2_. Biological n = 5 for G1100; n = 4 for G1096 and G1102; n = 2 for G1103; and n = 3 for all other strains shown. These full-period peak rates underlie the sensitivity summaries in the supplementary Results; the early-response rates and ratios in Figures 1 and 2 use a separate window search.

**Figure S9:**
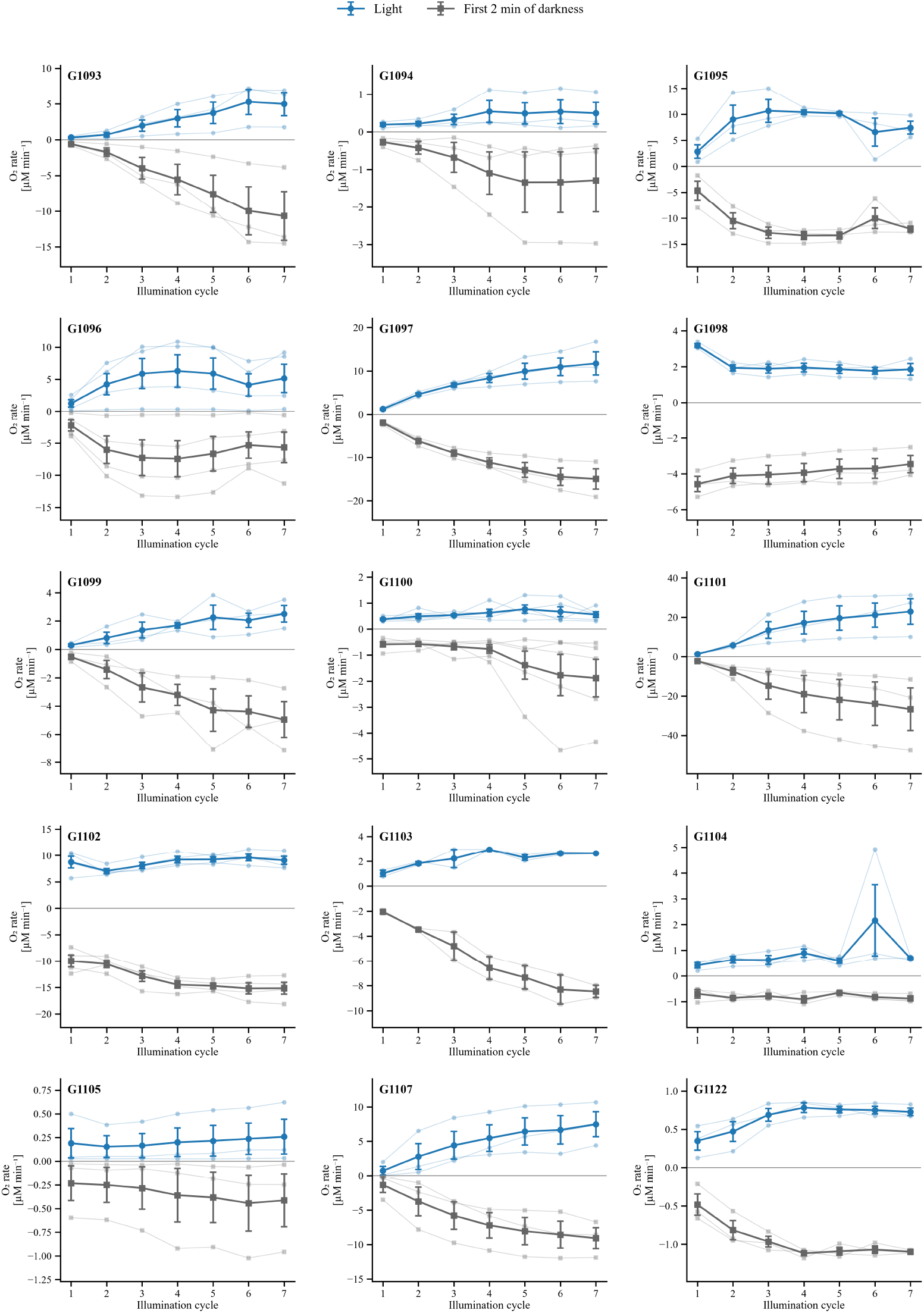
O_2_ rates through seven repeated light–dark cycles. Panels show 15 strains and 48 independently grown cultures, at each strain’s own constant irradiance (Table S3). Blue circles are light rates, and gray squares are rates during the first 2 min of the following dark interval. Thin pale lines connect values from individual cultures; the prominent lines and error bars show arithmetic means ± SEM. We selected rates from 30-s windows across each full 2-min analysis period using the signed selection rule in the Methods; positive and negative values mean net increases and decreases in dissolved O_2_. Biological n = 5 for G1100; n = 4 for G1096 and G1102; n = 2 for G1103; and n = 3 for all other strains shown. We kept all cycles; we did not assume that a later cycle had started below the operational O_2_ threshold. Panels follow numeric G-code order.

**Figure S10:**
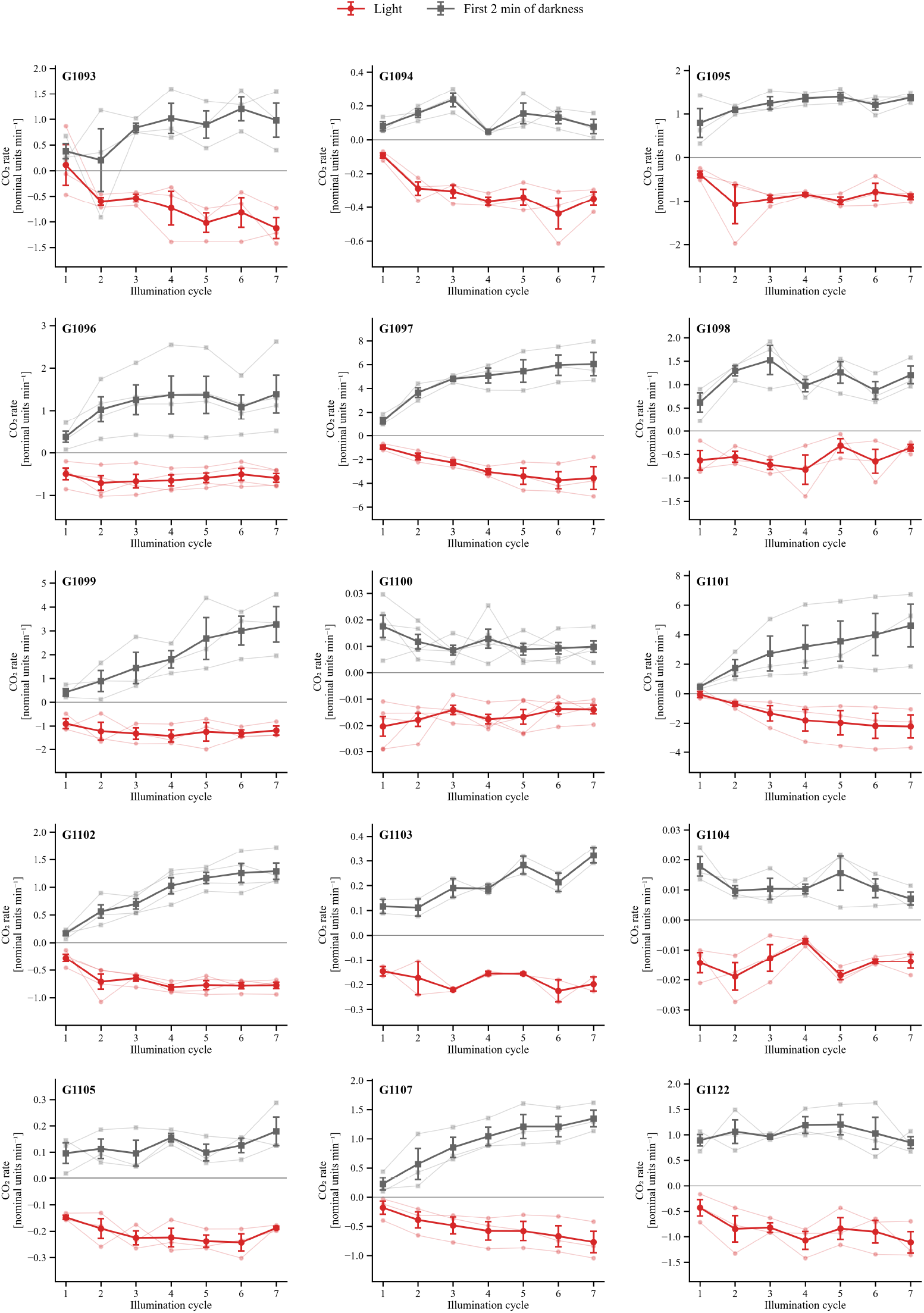
CO_2_ rates through seven repeated light–dark cycles. Panels show 15 strains and 48 independently grown cultures, at each strain’s own constant irradiance (Table S3). Red circles are light rates, and gray squares are rates during the first 2 min of the following dark interval. Thin pale lines connect values from individual cultures; the prominent lines and error bars show arithmetic means ± SEM. We obtained 30-s rates across each full 2-min analysis period, giving priority to negative slopes in light and positive slopes in darkness, as described in the Methods. Negative and positive values mean net decreases and increases in the CO_2_ signal. Biological n = 5 for G1100; n = 4 for G1096 and G1102; n = 2 for G1103; and n = 3 for all other strains shown. Panels follow numeric G-code order. CO_2_ axes use nominal signal units and nominal units min⁻¹ on the recorded conversion scale, not independently validated absolute concentrations or fluxes.

## Supplementary Methods

### TAP formulation

Per litre of TAP medium, the Chlamydomonas Resource Center’s reference recipe calls for: 20 mM Tris, 0.400 g NH₄Cl, 0.100 g MgSO₄·7H_2_O, 0.050 g CaCl_2_·2H_2_O, 0.108 g K_2_HPO₄, 0.056 g KH_2_PO₄, 1 mL Hutner’s trace-element stock, and 1 mL glacial acetic acid, adjusted to pH 7.0.

Per litre, the Hutner stock contains: 50 g disodium EDTA, 22 g ZnSO₄·7H_2_O, 11.4 g H₃BO₃, 5.06 g MnCl_2_·4H_2_O, 1.61 g CoCl_2_·6H_2_O, 1.57 g CuSO₄·5H_2_O, 1.10 g (NH₄)₆Mo₇O_2_₄·4H_2_O, and 4.99 g FeSO₄·7H_2_O. The reference source gives the stock-preparation and pH-adjustment procedure.

Recipe sources: Chlamydomonas Resource Center, “TAP and Tris minimal” (https://www.chlamycollection.org/methods/media-recipes/tap-and-tris-minimal/) and “Hutner’s trace elements” (https://www.chlamycollection.org/methods/media-recipes/hutners-trace-elements/). The TAP recipe is attributed to Gorman and Levine (1965), doi:10.1073/pnas.54.6.1665.

### Vitamin stock

The vitamin stock contained 100 mg thiamine hydrochloride, 1.5 mg biotin, 100 mg nicotinic acid, 1 mg vitamin B₁_2_ and 10 mg p-aminobenzoic acid in double-distilled water to 100 mL, sterilized by filtration. This is the shared stock composition documented in the companion characterization. The cultivation atmosphere for the present MIMS cultures was ambient air; the companion FireSting CO_2_-enrichment regime was not applied here.

### DMSO chlorophyll calculation

We centrifuged a 1 mL sample for 1 min at 16,000 × g, resuspended it in 1 mL of DMSO, incubated it at 65 °C for 15–20 minutes, centrifuged it again for 1 min at 16,000 × g, and measured its absorbance using the chlorophyll equation below:

Chlorophyll a = 12.19(A₆₆₅ − A₇₃₀) − 3.45(A₆₄₉ − A₇₃₀)

Chlorophyll b = 21.99(A₆₄₉ − A₇₃₀) − 5.32(A₆₆₅ − A₇₃₀)

Total chlorophyll = chlorophyll a + chlorophyll b (µg mL⁻¹).

A₆₆₅, A₆₄₉ and A₇₃₀ denote extract absorbances at 665, 649 and 730 nm; A₇₃₀ provides the baseline correction. The equations and extraction procedure follow Wellburn (1994), as in the companion characterization manuscript.

Reference: Wellburn, A. R. (1994). The spectral determination of chlorophylls a and b, as well as total carotenoids, using various solvents with spectrophotometers of different resolution. Journal of Plant Physiology 144: 307–313. doi:10.1016/S0176-1617(11)81192-2.

### Rate-window quality checks

Candidate 30-s windows started either at measured sample times or at the boundaries of the analysis period, and stayed within the 120-s analysis period. Each fit needed at least 10 measurements spanning at least 27 s, with no gap larger than 2.5 times the median sampling interval for that gas channel. These rules stop sparse or patchy sampling from producing a misleading 30-s rate. For each accepted window, we kept the regression slope, the coefficient of determination, and the residual error. We picked the peak using the direction rule from the main Methods, without imposing any R² threshold. Where possible, we also calculated fixed-first-30-s and full-120-s regressions for each phase.

For Figures 1 and 2 and Figures S3 and S4 and the ratios in Figure S2, the early-response rate calculation first located a raw extremum within the first 72 s of each phase, then tested 30-s fits starting 0–18 s later, in 0.6-s steps. For an expected rise we used a minimum and the most positive fitted slope; for an expected decline, a maximum and the most negative slope. H_2_/O_2_ were expected to rise in light and decline in darkness, and CO_2_ the reverse. We excluded samples at the exact phase endpoint from the preceding phase. The same minimum point count, span, and gap checks applied here too. We required all four first/late phase fits for a paired summary; we averaged the first and last-three rates within each culture, then across cultures with equal weight.

For the display alignment in Figure 1 and Figures S3 and S4, we looked for a local minimum before a rise (H_2_/O_2_) or a maximum before a fall (CO_2_), searching between 30 s before and 90 s after the documented light onset. We used a running median only to detect this origin point; we did not smooth the displayed or fitted concentrations themselves. We screened candidate changes against three times a robust noise estimate derived from concentration differences. When no qualifying origin was found, we used the measured point closest to the documented onset instead. We subtracted the selected time and concentration separately for each exposure, without stretching time or rescaling amplitude. Traces were interpolated onto a 1-s grid only within their measured range, leaving missing tails blank. We calculated the means for cycles 5–7 within each culture before averaging across cultures. Because each trace has its own time shift, the plotted 2-min/3-min background is only schematic and cannot mark each recording’s exact light-to-dark transition.

### First-to-late summaries

For each culture i, we defined the early-response late mean as Lᵢ = (rᵢ,₅ + rᵢ,₆ + rᵢ,₇)/3, and the paired difference as Δᵢ = rᵢ,₁ − Lᵢ. Figure 2 reports the arithmetic means of rᵢ,₁ and Lᵢ across cultures, their sample SDs (denominator n − 1), and the ratio of those two means. Table S4 reports the individual culture values, including the within-culture ratio rᵢ,₁/Lᵢ. Note that averaging those individual ratios is not the same as the ratio of means plotted in Figure 2. All 48 included cultures had complete paired H_2_ estimates, with positive first and later mean rates.

### Cross gas comparisons

For each of the 48 cultures, O_2_ and CO_2_ first-light rates and means of light rates from cycles 5– 7 were taken from the same early-response 30-s analysis, matched by experimental group and culture identifier to the H_2_ data (Tables S4, S6 and S7). The same selection algorithm was used for all gases, but windows were selected independently and were not necessarily simultaneous. Culture ratios were defined only when both rates were positive for H_2_/O_2_ or both were negative for CO_2_; zero or sign-changing endpoints were not assigned a ratio. Each strain’s ratio divided its mean first rate by its mean later rate, including all matched cultures in both means. This endpoint differs from the mean of culture ratios. Across strains, H_2_ ratios were compared with O_2_ and CO_2_ ratios by Spearman correlation (15 and 13 strains, respectively; Table S8). Two-sided Monte Carlo permutation p values used 999,999 random permutations, seed 20261025, and the correction (b + 1)/(B + 1), where b counts permuted absolute correlations at least as large as the observed value. Holm adjustment was applied across these two comparisons. Pearson correlations of log ratios were secondary sensitivity analyses with the same permutation procedure. Small sample sizes, selection of sign-consistent ratios, near-zero denominators and confounded assay conditions limit interpretation.

### Rate window sensitivity and coverage

The analysis used the calibrated per-phase rate estimates without refitting or changing the primary early-response endpoint. Full-period peak-30-s, fixed-first-30-s and whole-120-s estimates were taken from the same per-phase analysis. Primary early-response light rates were verified against all 48 unrounded paired culture records. For each endpoint, the first-light value was paired with the arithmetic mean of cycles 5–7 within the same culture, and strain means weighted cultures equally. The ratio of those two strain means was used consistently across endpoints, isolating the window definition from a change between ratio-of-means and median-of-culture-ratios summaries. Ratios were not assigned when a mean was nonpositive. All 48 cultures had complete whole-120-s and full-period peak estimates. Fixed-first-30-s estimates required eligible fits in all four relevant phases; 17 of 192 required phase windows failed the original ≥27-s observed-span criterion, leaving 36 complete culture pairs in 13 strains. The failed spans were 25.992–26.137 s. Those quality checks were retained. All four endpoints were also compared within the same 36-culture subset. Missing-window details and all signed culture values are included in the data files.

### O_2_ exclusion sensitivity definitions

The operational flag was a calibrated O_2_ median ≥10 µM during the 30 s before a documented sustained illumination. The initial-flag exclusion removed a complete culture if cycle 1 was flagged. The strict complete-case exclusion removed a culture if cycle 1 or any of cycles 5–7 was flagged; all three late cycles were retained for every remaining culture. These gave 45 and 44 cultures, respectively. Two additional descriptive alternatives retained each culture’s unflagged late cycles, weighting the resulting within-culture late means equally across cultures: omitting flagged late cycles alone left 46 cultures with 137 late-cycle observations; additionally omitting initial-flag cultures left 45 cultures with 134 late observations. A culture with no eligible late cycles was omitted from that alternative. These sensitivities do not replace the primary cohort or establish that unflagged cultures were truly anoxic independently of calibration uncertainty.

### Paired empirical bootstrap

Within each strain, the observed first and later rates from a culture were resampled together as a pair. All nⁿ ordered samples of n cultures with replacement were enumerated, giving 4–3125 ordered resamples for n = 2–5. For every resample, the mean first-minus-later difference and the ratio of first and later means were computed. Reported bounds are the 2.5th and 97.5th percentiles of the exact empirical-bootstrap distribution, using the inverse empirical cumulative distribution function. Enumeration eliminates Monte Carlo error but does not correct the limited information in the original sample. There are only three distinct culture-count vectors for n = 2 and ten for n = 3; resulting intervals are discrete, may have poor population coverage, and are used descriptively rather than as confirmatory significance tests. Leave-one-culture-out ratio ranges are also supplied in the data files. Resampling does not incorporate calibration uncertainty, gas withdrawal or instrument, irradiance and acquisition-date confounding.

### Unequal variance pairwise analysis

Games–Howell comparisons used the natural logarithm of each culture’s primary first-to-later H_2_ ratio. For each pair of strains, the standard error was √(sᵢ²/nᵢ + sⱼ²/nⱼ), with Welch– Satterthwaite degrees of freedom. The studentized-range statistic was √2 times the absolute Welch t statistic, using the family of 15 strains. All 105 pairwise p values were calculated; the 16 below 0.05 are shown in Table S13 and the full family is supplied in the data files. Studentized-range probabilities were evaluated by deterministic quadrature at 180 and 260 nodes in each dimension, with maximum absolute difference <4 × 10⁻¹⁴. Games–Howell avoids the common-variance assumption of pooled-error Tukey–Kramer comparisons, but the small samples and resulting degrees of freedom still limit inference. Differences of mean log culture ratios correspond to comparisons of geometric mean culture ratios and are distinct from the primary ratios of strain means. No pairwise non-detection is interpreted as equivalence.

### Sampling and fit diagnostics

Within-recording median sampling intervals in each of the H_2_, O_2_ and CO_2_ channels ranged from 2.124 to 2.599 s. This characterizes sampling cadence, not membrane or instrument response time. Primary selected H_2_ light fits began 0.001–39.700 s after the first sustained onset (median 16.061 s) and 6.444–78.247 s after late sustained onsets (median 26.244 s). The selected 30-s windows therefore ended before 120 s in every case. Their R² medians were 0.997 initially and 0.968 later; 15 of 144 late fits, and none of 48 initial fits, had R² <0.5. No additional R² filter was imposed. High R² is not validation of absolute calibration or exclusion of physical loss. In the fitting implementation, intervals are half-open at the documented phase endpoint, so an observation at exactly 120 s is excluded from the light interval.

## Supplementary Results

### Sensitivity to the rate summary

Using the full-period 30-s window search behind Figure S8, 43 of 48 cultures had a first-to-later H_2_ ratio above 1, and 14 of 15 strain median ratios exceeded 1. The largest medians were G1095 at 1110 µmol photons m⁻² s⁻¹ (14.13, range 12.94–20.23, n = 3), G1107 at 722 µmol photons m⁻² s⁻¹ (10.26, range 9.59–21.20, n = 3), and G1096 at 1110 µmol photons m⁻² s⁻¹ (8.81, range 1.55–17.64, n = 4). All ten cultures in these three strains kept a higher first than later rate when we used regressions across the complete 2-min light interval. G1103 had a median full-period ratio close to 1 (1.02, range 0.92–1.11, n = 2), and G1122 was the only strain with a median below 1 (0.71, n = 3).

The matching early-response ratios of group means, shown in Figures 1 and 2, were 17.48, 13.15 and 4.72 for G1095, G1107 and G1096. These two analyses differ in both the window rule and the summary statistic used, so their numerical differences cannot be attributed to window selection alone. That both analyses agree on the direction of the strongest responses supports a genuine first-light enhancement; but the exact size and ranking of the ratios still depend on which endpoint is used.

The full-period profiles also track the progression across individual cycles. Median H_2_ peak rates in cycles 1 and 7 were 6.50 and 0.58 µM min⁻¹ for G1095, 5.56 and 0.27 for G1107, and 17.66 and 3.09 for G1096. For G1098 these were 37.27 and 18.22 µM min⁻¹; for G1104, 9.29 and 6.03; and for G1103, 8.46 and 8.55. These cycle-specific values should not be used in place of the mean-first and mean-last-three early-response rates reported in Figure 2.

### Exploratory strain comparisons

After omitting G1095, G1096 and G1107, ordinary ANOVA gave F(11, 26) = 4.60, p = 0.000667, and Welch’s analysis gave F(11, 9.34) = 5.26, p = 0.00854. The full-data median-centered Brown–Forsythe test gave F(14, 33) = 2.20, p = 0.0310. The approximate one-way random-effects intraclass correlations were 0.728 for all strains and 0.533 after exclusion. These clustering summaries are distinct from fixed-effect η² = 0.802 and do not quantify a causal genetic contribution. Unequal-variance Games–Howell results are reported in Table S13, with the complete 105-pair family in Supplementary Data 1. With n = 2–5, even unequal-variance pairwise error control is uncertain; no non-detection is interpreted as equivalence.

### Late cycle behavior

The endpoint is an average of cycles 5–7 rather than a demonstrated plateau. Mean early-response H_2_ rates from cycle 5 to cycle 7 fell from 19.541 to 11.594 µM min⁻¹ in G1102, from 0.632 to 0.392 in G1107, from 1.340 to 0.939 in G1097, and from 12.531 to 9.437 in G1099. These descriptive changes were not formal stability tests, but show why a short dark interval alone cannot establish a stationary later response.

### Consistent aggregation across rate windows

With all 48 cultures, the ratio-of-strain-means ranges were 0.635–17.476 for the primary early-response endpoint, 0.660–16.895 for full-period peak-30-s slopes and 0.854–30.629 for whole-120-s slopes. Fourteen of fifteen strain means exceeded 1 for each endpoint. Primary and full-period peak methods agreed on the direction in every culture; whole-120-s values changed three culture-level directions (G1097 r02, G1103 r02 and G1122 r01) and gave 46 rather than 43 higher-first cultures. For G1095, G1096 and G1107, whole-period ratios of means were 22.874, 8.539 and 30.629, respectively, while all ten cultures retained the higher-first direction. These numerical changes are window effects under the same aggregation rule; the full-period median culture ratios reported above answer a different summary question.

In the fixed-first-30-s complete subset, all 36 cultures had higher first slopes, but 22 later slopes were nonpositive. Only 14 culture ratios and four strain ratios therefore met the positive-ratio definition. G1122’s mean first and later slopes were 3.403 and −0.627 µM min⁻¹, respectively; no positive first-to-later ratio was assigned. Within the identical 36-culture cohort, higher-first counts were 31 for early-response and full-period peak windows, 34 for whole-120-s slopes and 36 for fixed-first-30-s slopes. The corresponding numbers of strain ratios above 1 were 11/13, 11/13, 12/13 and 4/4 defined ratios, respectively; the remaining nine fixed-window strain ratios were undefined. Thus, onset-defined and response-selected windows describe different parts of the transient, and neither the exact ratio nor the inversion in G1122 is independent of that definition.

### O_2_ exclusion sensitivity

The initial-flag exclusion left 40/45 higher-first cultures and 13/15 strain means above 1. The strict initial-or-late complete-case exclusion left 39/44 higher-first cultures and 13/15 strain means above 1. G1097 changed from a primary ratio of 1.117 to 0.930 with the initial exclusion and to 0.714 under strict exclusion; its remaining sample sizes were two and one, respectively. G1093 changed from 1.674 to 1.422, with one culture remaining. G1095, G1096, G1107 and G1122 had no relevant O_2_ threshold flags, so their primary early-response values were unchanged. The available-late-cycle alternatives also yielded 13/15 strain means above 1, with G1097 at 0.897. These sensitivities support the broad predominance of higher first sustained-light rates while identifying G1097 as sensitive to the O_2_-based cohort definition.

### Uncertainty and pairwise sensitivity

Table S11 reports the primary paired means, mean differences and exact empirical-bootstrap percentile ranges for all 15 strains. G1096 illustrates ratio uncertainty: its primary ratio was 4.716, with an empirical-bootstrap range of 1.941–16.720 and a leave-one-out range of 3.227– 8.626. G1103’s two cultures give particularly sparse resampling support and cannot establish equivalence near a ratio of 1. Games–Howell yielded 16/105 adjusted p values below 0.05; all involved G1095, G1107 or G1122 and none involved G1096 or compared two of the other eleven strains. These unequal-variance comparisons reduce the 27 pooled-error Tukey–Kramer detections and reinforce that the available data do not support a complete strain ranking. Pairwise normal-model assumptions, small n and assay confounding remain limitations.

### Cross gas relationships

Matched estimates were available for all 48 cultures. O_2_ culture ratios were defined for 47 cultures; G1104 r01 had a negative first rate and a positive later rate. CO_2_ ratios were defined for 42 cultures; six cultures changed from a positive first slope to a negative later slope (Table S7). At the strain-mean level, all 15 O_2_ ratios were defined, whereas G1093 and G1094 were excluded from the CO_2_ ratio correlation because their mean slopes changed sign. The H_2_ ratio showed no detectable monotonic association with the O_2_ ratio (Spearman ρ = 0.007, n = 15, permutation p = 0.985, Holm-adjusted p = 0.985) or with the sign-consistent CO_2_ ratio (ρ = 0.363, n = 13, p = 0.223, Holm-adjusted p = 0.447). Secondary log-ratio Pearson correlations were −0.339 for O_2_ (p = 0.218) and 0.020 for CO_2_ (p = 0.949). These results do not provide evidence for a common cross-strain coupling of these endpoints, but their limited precision does not establish that physiological coupling is absent.

**Table S4:** Paired H_2_ rates for each independent culture behind Figures 1 and 2, in µM min⁻¹, using the early-response 30-s procedure. “Later” is the mean of cycles 5–7 within that culture; “difference” is first minus later. The final column is the within-culture ratio, which is different from the ratio of strain means plotted in Figure 2. Values are rounded to three decimals for display, but the calculations themselves used unrounded estimates. Table S3 gives the irradiances and strain sample counts.

| Strain | Culture | First | Later | Difference | Ratio |
| --- | --- | --- | --- | --- | --- |
| G1093 | r01 | 40.462 | 28.450 | 12.011 | 1.422 |
| G1093 | r02 | 66.022 | 39.362 | 26.660 | 1.677 |
| G1093 | r03 | 63.434 | 33.720 | 29.715 | 1.881 |
| G1094 | r01 | 37.555 | 19.179 | 18.376 | 1.958 |
| G1094 | r02 | 24.234 | 10.794 | 13.440 | 2.245 |
| G1094 | r03 | 57.666 | 30.078 | 27.588 | 1.917 |
| G1095 | r01 | 18.841 | 0.889 | 17.952 | 21.184 |
| G1095 | r02 | 5.804 | 0.395 | 5.409 | 14.680 |
| G1095 | r03 | 6.531 | 0.499 | 6.032 | 13.083 |
| G1096 | r01 | 21.360 | 4.982 | 16.377 | 4.287 |
| G1096 | r02 | 30.630 | 1.652 | 28.978 | 18.539 |
| G1096 | r03 | 14.026 | 1.019 | 13.007 | 13.769 |
| G1096 | r04 | 14.102 | 9.335 | 4.767 | 1.511 |
| G1097 | r01 | 1.366 | 0.798 | 0.568 | 1.712 |
| G1097 | r02 | 1.234 | 1.730 | -0.495 | 0.714 |
| G1097 | r03 | 1.121 | 0.803 | 0.318 | 1.397 |
| G1098 | r01 | 34.361 | 13.660 | 20.701 | 2.515 |
| G1098 | r02 | 37.464 | 19.027 | 18.437 | 1.969 |
| G1098 | r03 | 44.923 | 25.752 | 19.171 | 1.744 |
| G1099 | r01 | 23.336 | 12.938 | 10.398 | 1.804 |
| G1099 | r02 | 22.309 | 9.581 | 12.727 | 2.328 |
| G1099 | r03 | 21.565 | 9.847 | 11.718 | 2.190 |
| G1100 | r01 | 26.099 | 5.749 | 20.350 | 4.540 |
| G1100 | r02 | 8.317 | 5.262 | 3.055 | 1.581 |
| G1100 | r03 | 10.342 | 5.599 | 4.743 | 1.847 |
| G1100 | r04 | 12.319 | 4.340 | 7.979 | 2.839 |
| G1100 | r05 | 10.571 | 9.546 | 1.025 | 1.107 |
| G1101 | r01 | 4.700 | 3.890 | 0.810 | 1.208 |
| G1101 | r02 | 0.627 | 0.387 | 0.240 | 1.620 |
| G1101 | r03 | 0.684 | 0.145 | 0.539 | 4.711 |
| G1102 | r01 | 59.806 | 19.763 | 40.044 | 3.026 |
| G1102 | r02 | 48.099 | 22.188 | 25.912 | 2.168 |
| G1102 | r03 | 57.385 | 14.244 | 43.141 | 4.029 |
| G1102 | r04 | 32.975 | 5.362 | 27.614 | 6.150 |
| G1103 | r01 | 8.111 | 7.229 | 0.882 | 1.122 |
| G1103 | r02 | 8.818 | 9.510 | -0.692 | 0.927 |
| G1104 | r01 | 5.412 | 2.729 | 2.682 | 1.983 |
| G1104 | r02 | 9.645 | 5.972 | 3.673 | 1.615 |
| G1104 | r03 | 9.294 | 5.158 | 4.136 | 1.802 |
| G1105 | r01 | 16.144 | 6.135 | 10.009 | 2.631 |
| G1105 | r02 | 7.680 | 3.356 | 4.324 | 2.288 |
| G1105 | r03 | 12.177 | 5.846 | 6.331 | 2.083 |
| G1107 | r01 | 4.501 | 0.453 | 4.048 | 9.934 |
| G1107 | r02 | 5.544 | 0.557 | 4.987 | 9.958 |
| G1107 | r03 | 9.227 | 0.456 | 8.770 | 20.228 |
| G1122 | r01 | 13.371 | 15.962 | -2.591 | 0.838 |
| G1122 | r02 | 5.511 | 13.790 | -8.279 | 0.400 |
| G1122 | r03 | 7.509 | 11.837 | -4.328 | 0.634 |

**Table S5:** Pre-illumination O_2_ medians at or above 10 µM in the selected seven-cycle dataset. Each value is the median over the 30 s immediately before the documented light onset, using the original calibrated O_2_ signal. All 19 flagged starts are listed: three of the 48 initial starts, and 16 of the 288 later starts, including seven of the 144 starts in cycles 5–7. The initial flags disagree with our original anoxia assessment at acquisition time, and we kept them as calibration-dependent discrepancies. This classification does not depend on the H_2_ slope-selection method. Culture codes correspond to Table S4.

| Strain | Culture | Cycle | Pre-light O<sub>2</sub> (μM) |
| --- | --- | --- | --- |
| G1093 | r02 | 1 | 12.421 |
| G1093 | r02 | 2 | 11.885 |
| G1093 | r02 | 3 | 11.638 |
| G1093 | r02 | 4 | 11.565 |
| G1093 | r02 | 5 | 11.196 |
| G1093 | r02 | 6 | 11.046 |
| G1093 | r02 | 7 | 10.993 |
| G1093 | r03 | 1 | 10.791 |
| G1093 | r03 | 2 | 10.389 |
| G1093 | r03 | 3 | 10.283 |
| G1093 | r03 | 4 | 10.088 |
| G1097 | r01 | 1 | 11.136 |
| G1097 | r01 | 2 | 10.871 |
| G1097 | r01 | 3 | 10.991 |
| G1097 | r01 | 4 | 11.204 |
| G1097 | r01 | 5 | 11.500 |
| G1097 | r01 | 6 | 11.824 |
| G1097 | r01 | 7 | 12.329 |
| G1097 | r03 | 7 | 10.901 |

**Table S6:** Paired O_2_ light-period rates for all 48 independent cultures, using the same early-response selection algorithm as H_2_. Later is the mean of cycles 5–7 within the culture. H_2_ ratio is repeated from Table S4 for matching. O_2_ rates are µM min⁻¹. An em dash marks a sign-inconsistent ratio; first and later rates remain visible. No detection-limit threshold was imposed.

| Strain | Culture | H<sub>2</sub> ratio | O<sub>2</sub> first | O<sub>2</sub> later | O<sub>2</sub> ratio |
| --- | --- | --- | --- | --- | --- |
| G1093 | r01 | 1.422 | 0.231 | 0.616 | 0.375 |
| G1093 | r02 | 1.677 | 0.486 | 2.533 | 0.192 |
| G1093 | r03 | 1.881 | 0.192 | 1.281 | 0.150 |
| G1094 | r01 | 1.958 | 0.097 | 0.121 | 0.802 |
| G1094 | r02 | 2.245 | 0.265 | 0.291 | 0.911 |
| G1094 | r03 | 1.917 | 0.192 | 0.287 | 0.668 |
| G1095 | r01 | 21.184 | 0.981 | 2.405 | 0.408 |
| G1095 | r02 | 14.680 | 0.110 | 4.496 | 0.024 |
| G1095 | r03 | 13.083 | 0.251 | 6.952 | 0.036 |
| G1096 | r01 | 4.287 | 0.475 | 1.302 | 0.365 |
| G1096 | r02 | 18.539 | 1.955 | 4.129 | 0.473 |
| G1096 | r03 | 13.769 | 0.429 | 5.762 | 0.075 |
| G1096 | r04 | 1.511 | 0.088 | 0.236 | 0.373 |
| G1097 | r01 | 1.712 | 0.352 | 4.967 | 0.071 |
| G1097 | r02 | 0.714 | 0.489 | 4.197 | 0.117 |
| G1097 | r03 | 1.397 | 0.434 | 7.001 | 0.062 |
| G1098 | r01 | 2.515 | 3.080 | 0.845 | 3.644 |
| G1098 | r02 | 1.969 | 3.130 | 1.240 | 2.525 |
| G1098 | r03 | 1.744 | 3.420 | 0.650 | 5.265 |
| G1099 | r01 | 1.804 | 0.310 | 0.749 | 0.414 |
| G1099 | r02 | 2.328 | 0.222 | 0.308 | 0.720 |
| G1099 | r03 | 2.190 | 0.147 | 0.410 | 0.360 |
| G1100 | r01 | 4.540 | 0.274 | 0.359 | 0.765 |
| G1100 | r02 | 1.581 | 0.276 | 0.312 | 0.886 |
| G1100 | r03 | 1.847 | 0.220 | 0.444 | 0.496 |
| G1100 | r04 | 2.839 | 0.205 | 0.460 | 0.446 |
| G1100 | r05 | 1.107 | 0.507 | 0.563 | 0.901 |
| G1101 | r01 | 1.208 | 0.742 | 7.681 | 0.097 |
| G1101 | r02 | 1.620 | 0.436 | 2.617 | 0.167 |
| G1101 | r03 | 4.711 | 0.570 | 9.569 | 0.060 |
| G1102 | r01 | 3.026 | 9.770 | 6.099 | 1.602 |
| G1102 | r02 | 2.168 | 6.506 | 7.579 | 0.858 |
| G1102 | r03 | 4.029 | 8.448 | 7.629 | 1.107 |
| G1102 | r04 | 6.150 | 3.941 | 6.765 | 0.583 |
| G1103 | r01 | 1.122 | 0.388 | 0.435 | 0.892 |
| G1103 | r02 | 0.927 | 0.544 | 0.612 | 0.888 |
| G1104 | r01 | 1.983 | −0.017 | 0.549 | — |
| G1104 | r02 | 1.615 | 0.524 | 1.105 | 0.475 |
| G1104 | r03 | 1.802 | 0.390 | 0.618 | 0.631 |
| G1105 | r01 | 2.631 | 0.404 | 0.492 | 0.822 |
| G1105 | r02 | 2.288 | 0.033 | 0.051 | 0.653 |
| G1105 | r03 | 2.083 | 0.026 | 0.027 | 0.977 |
| G1107 | r01 | 9.934 | 0.011 | 2.044 | 0.005 |
| G1107 | r02 | 9.958 | 0.010 | 3.237 | 0.003 |
| G1107 | r03 | 20.228 | 1.114 | 6.695 | 0.166 |
| G1122 | r01 | 0.838 | 0.337 | 0.466 | 0.723 |
| G1122 | r02 | 0.400 | 0.112 | 0.394 | 0.285 |
| G1122 | r03 | 0.634 | 0.217 | 0.331 | 0.655 |

**Table S7:** Paired CO_2_ light-period rates for all 48 independent cultures. Definitions and matching follow Table S6. CO_2_ rates use nominal signal units min⁻¹; absolute concentrations and fluxes are not independently validated. The first-to-later ratio is shown only when both slopes are negative. An em dash identifies six first-positive sign reversals.

| Strain | Culture | H<sub>2</sub> ratio | CO<sub>2</sub> first | CO<sub>2</sub> later | CO<sub>2</sub> ratio |
| --- | --- | --- | --- | --- | --- |
| G1093 | r01 | 1.422 | 0.329 | -0.396 | — |
| G1093 | r02 | 1.677 | 0.355 | -0.563 | — |
| G1093 | r03 | 1.881 | -0.441 | -0.947 | 0.466 |
| G1094 | r01 | 1.958 | 0.101 | -0.309 | — |
| G1094 | r02 | 2.245 | -0.069 | -0.202 | 0.340 |
| G1094 | r03 | 1.917 | 0.163 | -0.229 | — |
| G1095 | r01 | 21.184 | -0.481 | -0.393 | 1.223 |
| G1095 | r02 | 14.680 | -0.225 | -0.226 | 0.996 |
| G1095 | r03 | 13.083 | -0.268 | -0.227 | 1.181 |
| G1096 | r01 | 4.287 | -0.378 | -0.393 | 0.962 |
| G1096 | r02 | 18.539 | -0.855 | -0.698 | 1.225 |
| G1096 | r03 | 13.769 | -0.517 | -0.443 | 1.166 |
| G1096 | r04 | 1.511 | -0.203 | -0.177 | 1.150 |
| G1097 | r01 | 1.712 | -0.419 | -0.823 | 0.509 |
| G1097 | r02 | 0.714 | -0.813 | -0.662 | 1.228 |
| G1097 | r03 | 1.397 | -0.298 | -1.150 | 0.259 |
| G1098 | r01 | 2.515 | -0.842 | -0.375 | 2.247 |
| G1098 | r02 | 1.969 | -0.095 | -0.097 | 0.984 |
| G1098 | r03 | 1.744 | -0.607 | -0.262 | 2.311 |
| G1099 | r01 | 1.804 | -0.479 | -0.474 | 1.012 |
| G1099 | r02 | 2.328 | -1.135 | -1.047 | 1.084 |
| G1099 | r03 | 2.190 | -0.958 | -1.294 | 0.741 |
| G1100 | r01 | 4.540 | -0.013 | -0.015 | 0.855 |
| G1100 | r02 | 1.581 | -0.029 | -0.008 | 3.536 |
| G1100 | r03 | 1.847 | -0.003 | -0.016 | 0.190 |
| G1100 | r04 | 2.839 | -0.002 | -0.010 | 0.205 |
| G1100 | r05 | 1.107 | -0.013 | -0.007 | 1.880 |
| G1101 | r01 | 1.208 | 0.102 | -0.488 | — |
| G1101 | r02 | 1.620 | −0.136 | −0.196 | 0.694 |
| G1101 | r03 | 4.711 | −0.191 | −0.919 | 0.208 |
| G1102 | r01 | 3.026 | −0.086 | −0.655 | 0.131 |
| G1102 | r02 | 2.168 | −0.275 | −0.711 | 0.386 |
| G1102 | r03 | 4.029 | −0.459 | −0.650 | 0.706 |
| G1102 | r04 | 6.150 | −0.191 | −0.239 | 0.801 |
| G1103 | r01 | 1.122 | −0.044 | −0.149 | 0.294 |
| G1103 | r02 | 0.927 | −0.166 | −0.127 | 1.306 |
| G1104 | r01 | 1.983 | −0.008 | −0.016 | 0.483 |
| G1104 | r02 | 1.615 | −0.018 | −0.010 | 1.781 |
| G1104 | r03 | 1.802 | −0.010 | −0.011 | 0.932 |
| G1105 | r01 | 2.631 | −0.164 | −0.215 | 0.764 |
| G1105 | r02 | 2.288 | −0.132 | −0.070 | 1.876 |
| G1105 | r03 | 2.083 | −0.157 | −0.180 | 0.872 |
| G1107 | r01 | 9.934 | −0.069 | −0.307 | 0.223 |
| G1107 | r02 | 9.958 | −0.030 | −0.219 | 0.135 |
| G1107 | r03 | 20.228 | −0.063 | −0.334 | 0.187 |
| G1122 | r01 | 0.838 | 0.247 | −0.412 | — |
| G1122 | r02 | 0.400 | −0.850 | −0.768 | 1.107 |
| G1122 | r03 | 0.634 | −0.293 | −0.858 | 0.341 |

**Table S8:** Strain first-to-later ratios used in the cross-gas correlations. Each ratio divides the equally weighted mean first-light rate by the mean later-light rate, with all n matched cultures included in both means. Ratios are dimensionless; irradiance is µmol photons m⁻² s⁻¹. Em dashes mark CO_2_ sign-changing strain means. These are ratios of means, not averages of culture ratios.

| Strain | n | Irradiance | H<sub>2</sub> ratio | O<sub>2</sub> ratio | CO<sub>2</sub> ratio |
| --- | --- | --- | --- | --- | --- |
| G1093 | 3 | 1110 | 1.674 | 0.205 | — |
| G1094 | 3 | 722 | 1.989 | 0.792 | — |
| G1095 | 3 | 1110 | 17.476 | 0.097 | 1.151 |
| G1096 | 4 | 1110 | 4.716 | 0.258 | 1.142 |
| G1097 | 3 | 722 | 1.117 | 0.079 | 0.581 |
| G1098 | 3 | 722 | 1.998 | 3.522 | 2.104 |
| G1099 | 3 | 1110 | 2.077 | 0.463 | 0.914 |
| G1100 | 5 | 1110 | 2.218 | 0.694 | 1.080 |
| G1101 | 3 | 1110 | 1.359 | 0.088 | 0.141 |
| G1102 | 4 | 357 | 3.221 | 1.021 | 0.448 |
| G1103 | 2 | 722 | 1.011 | 0.890 | 0.759 |
| G1104 | 3 | 107 | 1.757 | 0.395 | 0.965 |
| G1105 | 3 | 203 | 2.347 | 0.814 | 0.974 |
| G1107 | 3 | 722 | 13.146 | 0.095 | 0.187 |
| G1122 | 3 | 52 | 0.635 | 0.560 | 0.440 |

Spearman correlations with H<sub>2</sub>: O<sub>2</sub>, $\rho = 0.007$ (n = 15; permutation p = 0.985; Holm-adjusted p = 0.985); CO<sub>2</sub>, $\rho = 0.363$ (n = 13; p = 0.223; Holm-adjusted p = 0.447). Windows were selected independently for each gas, so these analyses do not measure simultaneous electron partitioning.

**Table S9:** Collection records for the 17 MIMS strains. Columns and shared-strain records follow Table S1 of the companion oxygen characterization manuscript. Additional collection records are reported where available; unfilled fields are left blank. Sample dates refer to collection records rather than the MIMS assays.

| Index | Code | Identification | Algae-Base ID | Sample Region | Sample Location | Latitude | Longitude | Sample Date |
| --- | --- | --- | --- | --- | --- | --- | --- | --- |
| 1 | G1093 | *Chlorella ohadii* | 177095 | North-West Negev | Nitzana Dunes (Biological sand crust) | 30.933333 | 34.383333 | 2012 |
| 2 | G1094 | *Chlorella sorokiniana* | 51508 | Tel Aviv-Yafo | Yarkon River | 32.09623173 | 34.78216334 | 2024 |
| 3 | G1095 | *Micractinium tetrahymenae* | 176401 | Tel Aviv-Yafo | Yarkon River | 32.09623173 | 34.78216334 | 2024 |
| 4 | G1096 | *Desmodesmus abundans* | 42384 | Lake Kinneret | Kursi Beach | 34.492929 | 35.384136 | 2023 |
| 5 | G1097 | *Tetradismus obliquus* | 158061 | Negev | Ein Yorqe'am | 31.03166667 | 35.04138889 | 2024 |
| 6 | G1098 | *Chlamydomonas reinhardtii* CC124 |  |  | Reference culture |  |  |  |
| 7 | G1099 | *Chlorella* sp. | 43426 | Golan Heights | Shaavnia Reservoir | 32.89444444 | 35.78166667 | 2023 |
| 8 | G1100 | *Chlorella* sp. | 43426 | HaSharon | Hadera Stream Park | 32.46805556 | 34.90305556 | 2023 |
| 9 | G1101 | *Chlorolobion braunii* | 27677 | Negev | Ein Yorqe'am | 31.03166667 | 35.04138889 | 2024 |
| 10 | G1102 | *Acutodesmus obliquus* SAG 276-3d | 135513 | Italy | Castellamare | 40.706018 | 14.483042 | 1954 |
| 11 | G1103 | *Lobochlamys segnis* | 118596 | Golan Heights | El-Al Stream | 32.82861111 | 35.89888889 | 2023 |
| 12 | G1104 | *Monoraphidium cf. dybowskii* | 131950 | Yarkon River | Outflow | 32.09623173 | 34.78216334 | 2024 |
| 13 | G1105 | *Micractinium* sp. | 43421 | HaSharon | Hadera Stream Park | 32.46805556 | 34.90305556 | 2023 |
| 14 | G1106 | *Stichococcus bacillaris* | 1044 | Northern District | Dardara Park |  |  | 2023 |
| 15 | G1107 | *Desmodesmus abundans* | 42384 | Golan Heights | Winter puddle near Bental reservoir | 33.23722222 | 36.30527778 | 2023 |
| 16 | G1121 | *Chlorella lewinii* | 147234 | Modi'in | Modi'in | 31.535084 | 34.590826 | 2025 |
| 17 | G1122 | *Monoraphidium convolutum* | 27720 | Negev | Gev Zarhan | 30.83388889 | 34.95833333 | 2024 |

**Table S10:** H_2_ sensitivity to rate window with a consistent ratio of strain means. First and later means use equal culture weights. Later is the within-culture mean of cycles 5–7. Primary, full-period peak and whole-period estimates are complete in all 48 cultures. Fixed-first-30-s estimates require complete eligible fits for cycles 1,5,6,7, yielding 36 cultures in 13 strains; a dash in the ratio column denotes nonpositive mean first or later rate, or no complete pairs. Fixed first/later means are µM min−1 and retain their signs. The separate common-36-culture analysis removes coverage differences from comparisons of window definitions. The n column applies to the primary, peak-30-s and whole-120-s endpoints.

| Strain | n | Early 30 s ratio | Peak 30 s ratio | Whole 120 s ratio | n fixed 30 s | Fixed first/late means | Fixed 30 s ratio |
| --- | --- | --- | --- | --- | --- | --- | --- |
| G1093 | 3 | 1.674 | 1.692 | 2.563 | 3 | 34.192 / -1.795 | — |
| G1094 | 3 | 1.989 | 2.014 | 2.478 | 1 | 31.804 / -7.818 | — |
| G1095 | 3 | 17.476 | 16.895 | 22.874 | 3 | 8.080 / -0.067 | — |
| G1096 | 4 | 4.716 | 4.769 | 8.539 | 4 | 13.513 / 0.477 | 28.333 |
| G1097 | 3 | 1.117 | 1.195 | 1.679 | 3 | 0.614 / -0.070 | — |
| G1098 | 3 | 1.998 | 1.973 | 2.430 | 2 | 24.912 / -0.073 | — |
| G1099 | 3 | 2.077 | 2.069 | 2.367 | 3 | 11.596 / -0.804 | — |
| G1100 | 5 | 2.218 | 1.591 | 4.944 | 0 | — | — |
| G1101 | 3 | 1.359 | 1.373 | 5.699 | 3 | 1.839 / 0.205 | 8.949 |
| G1102 | 4 | 3.221 | 3.250 | 3.161 | 4 | 25.004 / 0.687 | 36.418 |
| G1103 | 2 | 1.011 | 1.005 | 1.126 | 1 | 6.774 / 0.680 | 9.965 |
| G1104 | 3 | 1.757 | 1.422 | 2.192 | 0 | — | — |
| G1105 | 3 | 2.347 | 2.362 | 3.172 | 3 | 6.611 / -0.942 | — |
| G1107 | 3 | 13.146 | 13.252 | 30.629 | 3 | 2.712 / -0.093 | — |
| G1122 | 3 | 0.635 | 0.660 | 0.854 | 3 | 3.403 / -0.627 | — |

**Table S11:**
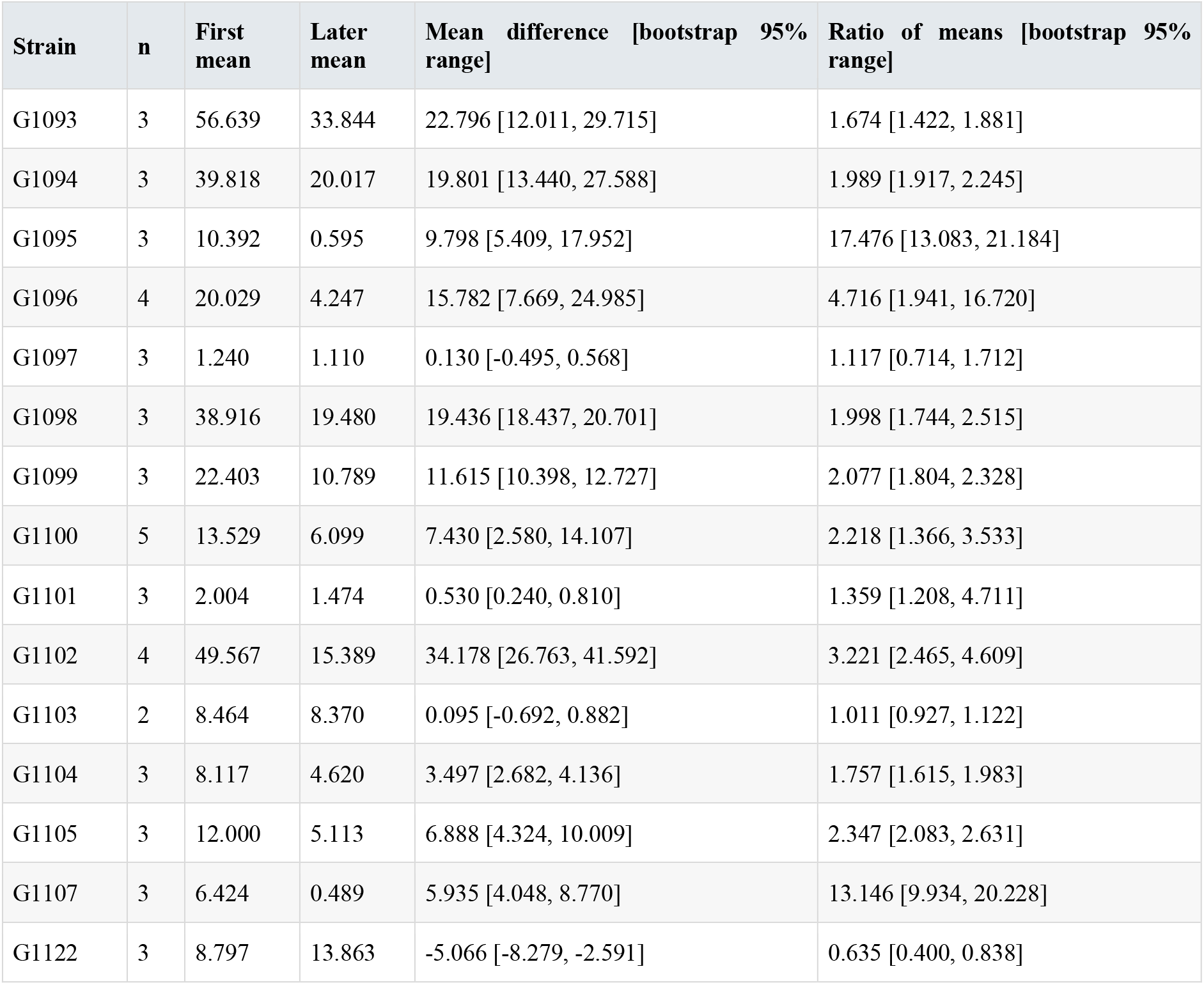
Paired empirical bootstrap uncertainty of the primary H_2_ endpoint. Rates and differences are µM min−1. First and later values from the same culture were resampled as pairs within each strain, enumerating all n^n ordered n-out-of-n resamples. Bounds are inverse-empirical-CDF 2.5th and 97.5th percentiles, with no Monte Carlo error. These are descriptive empirical-bootstrap intervals, not confirmatory tests or corrections for calibration or assay confounding. With n=2–5, support is sparse and coverage may be poor; G1103 has only three distinct count vectors.

**Table S12:** Sensitivity of primary H_2_ ratios to operational O_2_ threshold exclusions. A flag is a calibrated O_2_ median ≥10 µM during the 30 s before the relevant sustained illumination. Exclusions are sensitivity analyses, not changes to the primary cohort. Complete-case exclusion removes a culture if cycle 1 or any of cycles 5–7 is flagged and keeps all three late cycles for each remaining culture. Totals are 48, 45 and 44 cultures respectively; the strict sensitivity leaves n=1 in G1093 and G1097. The two available-cycle alternatives are reported in the machine-readable supplement. Absolute-calibration uncertainty remains.

| Strain | All cultures n / ratio | Exclude initial flags n / ratio | Exclude any initial or cycles 5–7 flags n / ratio |
| --- | --- | --- | --- |
| G1093 | 3 / 1.674 | 1 / 1.422 | 1 / 1.422 |
| G1094 | 3 / 1.989 | 3 / 1.989 | 3 / 1.989 |
| G1095 | 3 / 17.476 | 3 / 17.476 | 3 / 17.476 |
| G1096 | 4 / 4.716 | 4 / 4.716 | 4 / 4.716 |
| G1097 | 3 / 1.117 | 2 / 0.930 | 1 / 0.714 |
| G1098 | 3 / 1.998 | 3 / 1.998 | 3 / 1.998 |
| G1099 | 3 / 2.077 | 3 / 2.077 | 3 / 2.077 |
| G1100 | 5 / 2.218 | 5 / 2.218 | 5 / 2.218 |
| G1101 | 3 / 1.359 | 3 / 1.359 | 3 / 1.359 |
| G1102 | 4 / 3.221 | 4 / 3.221 | 4 / 3.221 |
| G1103 | 2 / 1.011 | 2 / 1.011 | 2 / 1.011 |
| G1104 | 3 / 1.757 | 3 / 1.757 | 3 / 1.757 |
| G1105 | 3 / 2.347 | 3 / 2.347 | 3 / 2.347 |
| G1107 | 3 / 13.146 | 3 / 13.146 | 3 / 13.146 |
| G1122 | 3 / 0.635 | 3 / 0.635 | 3 / 0.635 |

**Table S13:** Games–Howell comparisons of log culture-level H_2_ ratios with adjusted p < 0.05. All 105 pairs among 15 strains were examined using natural-log within-culture first/late H_2_ ratios, unpooled variances and Welch–Satterthwaite degrees of freedom; p values use the studentized-range distribution with 15 groups. Sixteen pairs met adjusted p<0.05; all involve G1095, G1107 or G1122, none involves G1096, and none compares two of the eleven middle strains. Ratios in this table compare geometric means of culture ratios, not the primary ratio of strain means. With n=2–5 and few pairwise degrees of freedom, these are exploratory sensitivity results, not a complete ranking or evidence of equivalence.

| Strain A | Strain B | n A / B | Geometric mean ratio A / B | Welch df | Games–Howell p |
| --- | --- | --- | --- | --- | --- |
| G1093 | G1095 | 3 / 3 | 0.103 | 3.14 | 0.00746 |
| G1094 | G1095 | 3 / 3 | 0.127 | 2.46 | 0.01867 |
| G1095 | G1097 | 3 / 3 | 13.359 | 3.11 | 0.03059 |
| G1095 | G1098 | 3 / 3 | 7.780 | 3.69 | 0.00726 |
| G1095 | G1099 | 3 / 3 | 7.620 | 3.04 | 0.01132 |
| G1095 | G1100 | 3 / 5 | 7.572 | 5.86 | 0.01007 |
| G1095 | G1102 | 3 / 4 | 4.471 | 4.81 | 0.04658 |
| G1095 | G1103 | 3 / 2 | 15.652 | 2.99 | 0.00586 |
| G1095 | G1104 | 3 / 3 | 8.901 | 2.65 | 0.01326 |
| G1095 | G1105 | 3 / 3 | 6.871 | 2.83 | 0.01558 |
| G1095 | G1122 | 3 / 3 | 26.757 | 3.50 | 0.00612 |
| G1097 | G1107 | 3 / 3 | 0.095 | 3.95 | 0.03885 |
| G1100 | G1107 | 5 / 3 | 0.167 | 5.44 | 0.04986 |
| G1102 | G1122 | 4 / 3 | 5.985 | 4.84 | 0.04191 |
| G1103 | G1107 | 2 / 3 | 0.081 | 2.57 | 0.03484 |
| G1107 | G1122 | 3 / 3 | 21.121 | 3.97 | 0.01047 |

## Notes

### Competing Interest Statement

The authors have declared no competing interest.

